# Human metabolites modulate antifungal efficacy and reveal creatinine-mediated antagonism of flucytosine

**DOI:** 10.64898/2026.06.12.731662

**Authors:** Johannes Hartl, Andrea Lehmann, Fiona Wahl, Markus Ralser

## Abstract

Clearing fungal infections can fail, even when the causative pathogen is susceptible to antifungal drugs in vitro. The reasons for this discrepancy often remain unclear. Here, we report that human metabolites affect antifungal drug efficacy. Studying *Saccharomyces cerevisiae,* we observed that human serum and common growth supplements alter the minimal inhibitory concentrations (MIC) across major antifungal classes, including fluconazole (FLC) and flucytosine (5-FC). To identify causal metabolites, we generated a library reflecting the physiological concentrations of 67 abundant blood metabolites. Assessing their impact on fungal fitness in the presence of antifungals revealed that creatinine, owing to its structural similarity to 5-FC, reduced the uptake and efficacy of 5-FC in *S. cerevisiae, Candida albicans,* and *Nakaseomyces glabratus*. Notably, 5-FC efficacy in urinary tract infections can be limited, and creatinine is highly concentrated in urine. We show that creatinine levels in urine increase the MIC of 5-FC in *N. glabratus* by approximately 100-fold, while reducing creatinine concentration in urine enhances 5-FC efficacy. Together, our findings highlight that drug interactions with creatinine and other human metabolites can strongly influence the activity of antifungals.

## INTRODUCTION

Fungal infections impose a major burden on human health, ranging from common mucosal and cutaneous diseases to invasive infections that remain associated with unacceptably high mortality despite antifungal therapy [1,2]. Indeed, latest estimates suggest that collectively up to 2.5 million people die annually from fungal infections [3]. Among these, bloodstream infections of yeasts such as *Candida albicans* and *Nakaseomyces glabaratus* are particularly problematic [1,3]. Accordingly, they have been classified as critical and high priority fungal pathogens by the World Health Organisation [4]. In parallel, a large proportion of the human population experience mucosal and superficial fungal infections, affecting the urogenital tract, as well as skin, nails and hair. While these infections are typically not life threatening, they still have a high medical burden, can cause significant discomfort, are a source of inflammation, and can be a comorbidity to other indications [5–7].

Antifungal treatment is currently based on only four major antifungal drug classes [8,9], and the arsenal is difficult to expand, with the genetic similarity of fungi to humans [10]. Major clinical antifungals are i) azoles targeting biosynthesis of the fungal sterol, ergosterol, ii) echinocandins, inhibiting biosynthesis of the cell wall component beta-glucan, iii) polyenes, binding ergosterol and increasing membrane permeability, and iv) pyrimidine analogs, inhibiting DNA and RNA metabolism and synthesis [11].

Fortunately, despite being on the rise, intrinsic drug resilience, such as genetically transmitted drug resistance to this limited arsenal of antifungals remains overall low [9,12–14]. However, despite the general susceptibility, pathogens are not always cleared by drugs *in vivo*: *[13,15,16]*. This situation can be caused by host factors such as immune status, fungal factors, or their interactions [2]). Further, divergent drug responses and efficacy may arise due to differences in pharmaco-kinetics and -dynamics [17,18], as well as differential drug responses in cell sub-populations, leading to fungal drug tolerance [9,19–21].

Although the role of the host and environmental factors in drug responses is widely accepted, the complexity of the potential interactions render this problem underinvestigated. Notably, for bacteria it has recently been shown that host niche can alter the efficacy of antimicrobials, and increase the minimal inhibitory concentration (MICs) of antimicrobials [22]. Also in yeasts, different environments can alter drug efficacy either directly, e.g. by changing pH, or indirectly, by altering fungal physiology that in turn affects fungal susceptibility to antifungal challenge [19,23,24]. A seminal example of the latter is the impact of carbon sources available to the pathogen [24,25]. For instance, switching from glucose to lactate-based media alters the cell wall architecture of *Candida albicans,* affecting fungal sensitivity to echinocandin or polyene drugs, as well as its interaction with the host immune system [24,26,27]. Similarly, *Cryptococcus neoformans’* susceptibility to the polyene Amphotericin B (AmB) is affected by glucose signaling, with higher glucose concentrations in the brain compared to e.g. lung altering the lipid profile to render a subpopulation of cells tolerant to the drug [20]. Lastly, drugs prescribed in conditions where fungal infections manifest as a comorbidity, also affect the efficacy of antifungals [28]. However, despite growing evidence that environmental and host-associated factors modulate antifungal efficacy, we often lack a systematic and causal understanding of how physiological biochemical interactions shape antifungal drug efficacy.

## RESULTS

### Metabolic environments affect the efficacy of common antifungals

Using liquid microbroth dilution assays (MDA), we started by determining the inhibitory concentrations for antifungals representing the key clinically used substance classes, namely an azole (fluconazole, FLC), an echinocandin (caspofungin, CSP), a polyene (amphotericin B, AmB), and an antimetabolite (flucytosine, 5-FC) in *S. cerevisiae* [11]. We tested these antifungals on glucose synthetic minimal (SM) media, SM supplemented with a mix of amino acids and nucleobases (synthetic complete, SC), a cell culture media RPMI-1640 (RPMI), and yeast extract and peptone with glucose (YPD) medium. Furthermore, to address the potential role of a physiological matrix, we supplemented SM also with 10% fetal calf serum (SM+FCS) (Fig. 1A). In order to exclude an effect of media pH differences, all growth media were buffered with MOPS at pH 6.75. The efficacy of each tested antifungal was influenced by the media composition, and each media and drug produced a specific pattern (Fig. 1A). For example, although antifungals were generally more effective in RPMI, CSP showed low effectiveness in this medium. In contrast, cells grown in YPD were more sensitive to CSP, while they were most resilient to 5-FC and FLC in this condition. As these differences were not explained by different growth rates alone (Fig. 1B), we speculated that they are explained by the media components. Here, the spread of 5-FC responses was of particular interest, as differences in its minimal inhibitory concentration were pronounced between environments (spanning ∼1.25 ng/ml to 640 ng/ml, Fig. 1A). 5-FC acts as a prodrug, which enters the cell via the pyrimidine-purine transporter Fcy2p and related permeases [29], where it is metabolized by the cytosine-deaminase Fcy1p to the chemotherapeutic agent flu-uracil, which after conversion by Fur1p is metabolized to inhibit thymidylate synthetase (Cdc21p) or incorporate into RNA, inhibiting protein synthesis [30] (Fig. 1C). A deletion strain of *fcy1* was thus resistant to 5-FC (Fig. 1D), and in line with its role as the major uptake system for 5-FC, *Δfcy2* was also largely insensitive to the drug (Fig. 1D).

**Figure 1.**
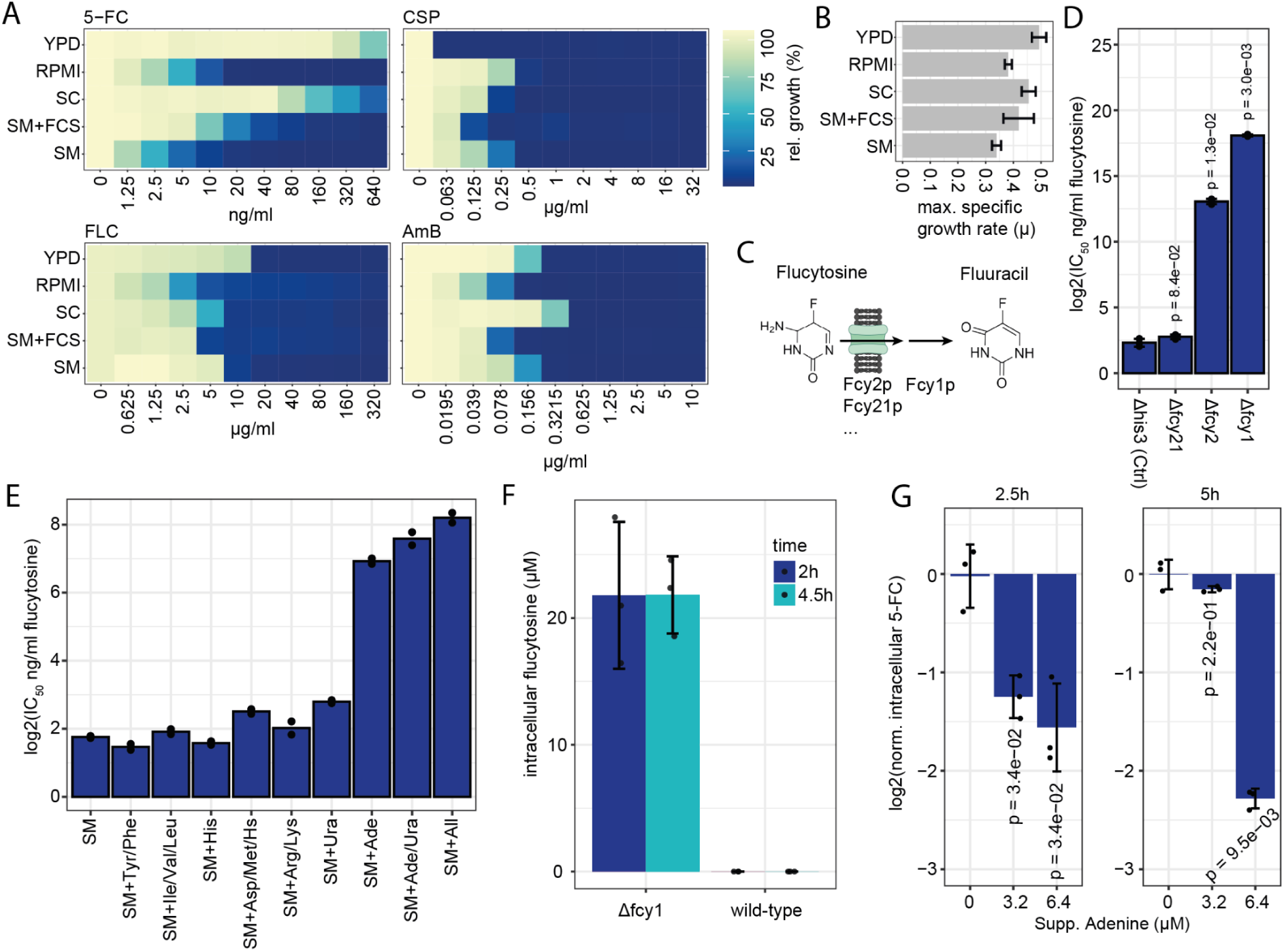
Growth media and metabolite supplements modulate antifungal susceptibility profiles, particularly for flucytosine. **(A)** *S. cerevisiae* was grown in commonly used media, and antifungal susceptibility was assessed by microbroth dilution assays with increasing antifungal concentrations. Relative growth after 24h is shown as OD_600_ in drug-treated wells normalized to the corresponding untreated control for each antifungal. Shown are mean values of n=3 replicates. **(B)** Corresponding maximum specific growth rates (μ) in each medium, calculated from the untreated growth experiments shown in (A). **(C)** Schematic representation of flucytosine uptake and metabolism, highlighting key genes involved in flucytosine uptake and deamination to the active compound fluuracil. **(D)** IC_50_ values for deletion strains of the key genes shown in (C) from the *S. cerevisiae* prototrophic knockout library, estimated by fitting dose–response curves to microbroth dilution data, here using the LL.4 model (drc R package). Shown are mean values of n=3 replicates +/- SD. Statistics based on t-tests comparing each deletion strain to the indicated control, with Holm correction for multiple testing. **(E)** IC_50_ values against flucytosine, determined by dose-response curves of microbroth dilution data in synthetic minimal (SM) medium, or supplemented with the indicated amino acid combinations (“drop-ins”). Adenine supplementation particularly reduced flucytosine efficacy. A prototrophic *S. cerevisiae* strain was used. “SM+All” denotes SM supplemented with the full combination of amino acids shown. Shown are mean values of n=2 replicates. **(F)** Intracellular flucytosine concentrations were quantified by LC–MS/MS in the Δfcy1 deletion strain and the corresponding control strain. Note that flucytosine was not detected in the wild-type strain. Shown are mean values of n=3 replicates +/- SD. **(G)** Supplementation of SM medium with adenine reduced intracellular flucytosine levels as estimated by LC-MS/MS in Δfcy1 cells after 2.5 and 5h of growth in a dose-dependent manner. Shown are mean values of n=3 replicates +/- SD. Statistics based on t-tests comparing each adenine concentration to the unsupplemented control, with Holm correction for multiple testing.

To identify the components responsible for the different 5-FC responses, we created several drop out media, on the basis of the SM and SC media recipes, respectively. We found that adenine, and to a lesser extent potentially uracil, influenced 5-FC’s effectiveness in inhibiting fungal growth (Fig. 1E). This is consistent with early work showing that adenine and related compounds antagonize 5-FC [31]. To confirm this, we set up a method for intracellular quantification of 5-FC. We established a rapid filtration-based workflow combining fast vacuum-assisted washing to quickly remove extracellular drug with subsequent hot ethanol extraction and LC–MS/MS analysis using an isotope-labeled internal standard. The drug 5-FC was reliably detected in *S. cerevisiae Δfcy1* strain exposed to the drug, but below the detection limit of the assay in wild-type (Fig. 1F). Importantly, adenine supplementation of *Δfcy1* cells exposed to 5-FC drastically reduced intracellular drug levels in concentration dependent manner (Fig. 1G).

Taken together, the efficacy of all major clinical antifungal classes on slowing growth in *S. cerevisiae* was dependent on the growth media. We studied 5-FC in more detail, and report that the supplementation of adenine decreases intracellular 5-FC levels and with it, the efficacy of this antifungal. Thus, at least in part, these effects are due to altered uptake or efflux of the drugs.

### Biochemical reconstitution of a human serum metabolome reveals its effect on 5-FC and FLC efficacy

Next, we tested if human serum (HS), as a representative matrix of a relative infection niche, would affect antifungal activities as well. Indeed, media supplementation with 10% HS reduced the MIC of AmB, FLC as well as CSP, and increased the MIC for 5-FC (Fig. 2A). We then asked if specific metabolites of the HS explain the effect. Serum includes a large variety of metabolites, lipids and proteins [32–34]. While the effects of small molecules on antifungals is understudied, the composition of the blood-metabolome is well characterized [32]. This allowed the re-creation of serum-like metabolite matrices [35,36]. Building upon the approach taken in these studies, we generated a concentration mimetic for the most abundant and accessible serum metabolites. Our focus was on biochemical supplements, and metabolites contained in yeast minimal media formulations, such as glucose, vitamins, etc. were not included. To assure that the selection and concentration estimates reflect current knowledge of the serum metabolome, we mined metabolomics datasets from healthy adult human serum in the human metabolome database [32]. We primarily selected metabolites that were abundant (>10µM), water-soluble, and commercially available, resulting in a synthetic collection of 67 metabolites, the human blood metabolite mixture (HBMM) (Fig. 2B, Supplementary Table 1, Methods). For most metabolites, abundance and identity closely matched previous reconstructions of the serum metabolome (Supplementary Fig. 1A, B) [35,36]. In total, 43 of supplemented metabolites were shared across all three studies. HBMM contained 19 additional compounds, whereas 11 predominantly low-abundance molecules reported previously were not included in our set (Supplementary Fig. 1A, B). The metabolites of our HBMM span a variety of functions and physicochemical properties, including amino acids and their derivatives, organic acids and carbohydrates (Fig. 2B). We obtained the compounds commercially and dissolved them as concentrated stock solutions to ultimately match the estimated blood concentration range of 0.01 to 5 mM (Supplementary Fig. 1A, Supplementary Table 1). On the basis of the full HBMM, we assessed its impact on yeast physiology. First, we performed a proteomic analysis of yeast cells with HBMM added to minimal medium. Adding the HBMM substantially altered proteome profiles, with ∼15% (221/3385) robustly quantified proteins showing differential abundance (Fig. 2C). KEGG pathway enrichment analysis indicated that proteins involved in amino acid biosynthesis, purine metabolism, and carbohydrate metabolism were among the most strongly affected (Supplementary Fig. 2A). These changes were partially explained by repression of biosynthesis pathways, e.g. for amino acids, which is in line with enrichment of *GCN4* and *LEU3* transcription factor motifs among differentially expressed genes (Supplementary Fig. 2B). These results support that yeast senses, uptakes, and metabolizes blood metabolites at physiological concentration.

**Figure 2.**
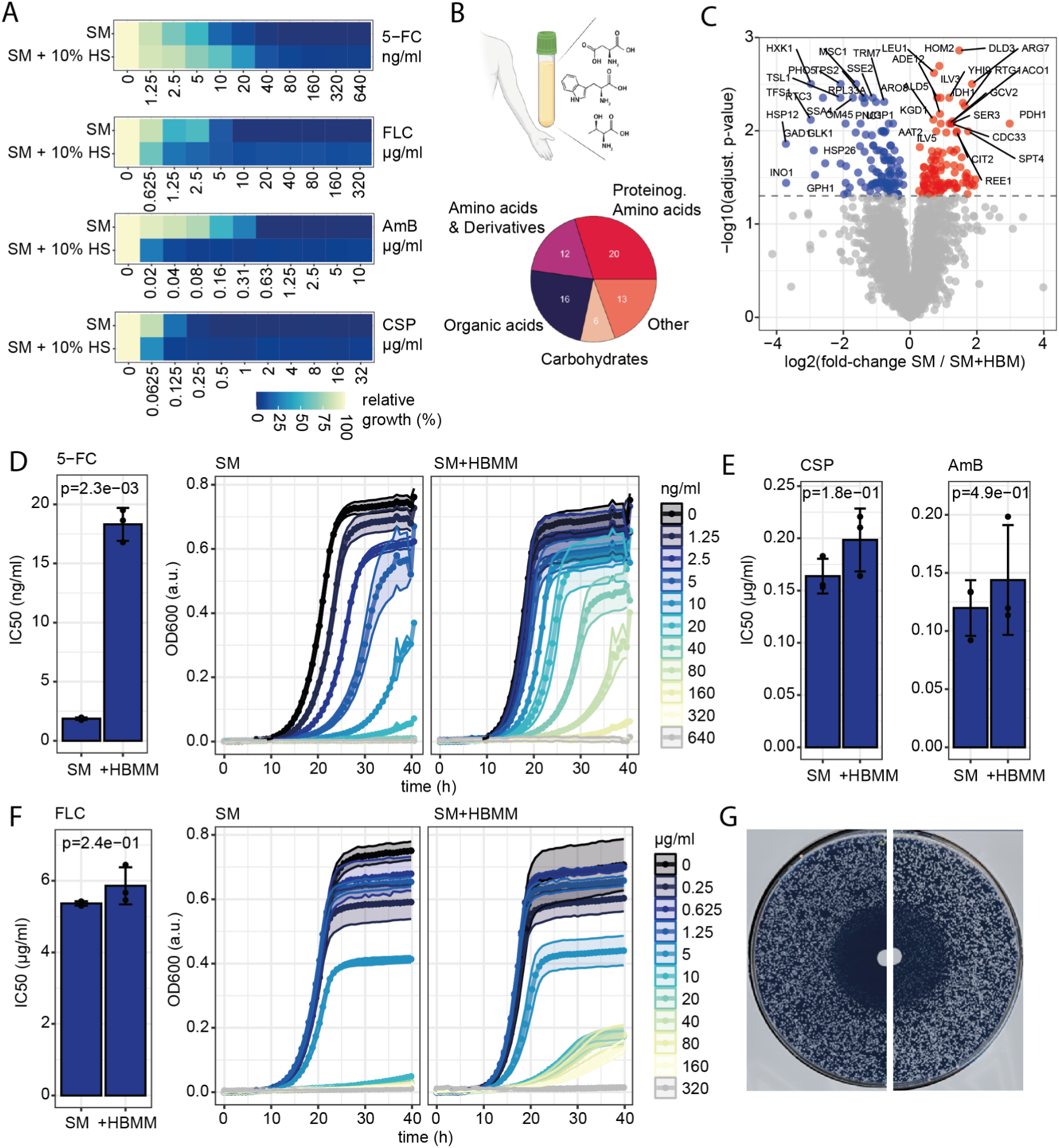
Human blood metabolites affect fungal susceptibility to flucytosine and fluconazole. **(A)** *S. cerevisiae* was grown in SM medium with or without 10% human serum, and antifungal susceptibility was assessed by microbroth dilution with increasing antifungal concentrations. Relative growth after 24 h is shown as OD_600_ in drug-treated wells normalized to the corresponding untreated control. Data based on mean values of n=3 replicates. **(B)** Schematic indicating the human blood metabolite mix (HBMM), a defined mixture of 67 metabolites abundant in human blood representing the indicated biochemical classes, and assembled at estimated physiological concentrations using the HMDB database. **(C)** Volcano plot of proteomics data with n=4 replicates per condition comparing *S. cerevisiae* grown in SM versus SM supplemented with the HBMM at physiological concentrations. Statistics were calculated using t-tests with Benjamini–Hochberg false discovery rate (FDR) correction. **(D)** Flucytosine sensitivity of *S. cerevisiae* was assessed by microbroth dilution in synthetic minimal (SM) medium or SM supplemented with HBMM. IC_50_ values were estimated from dose–response curves using OD_600_ after 24h (left), and corresponding growth curves are shown (right). Statistics were calculated using t-tests. Data from n=3 replicates, shown are mean values +/- SD. **(E)** IC_50_ values were determined as in (D) for caspofungin (left) and amphotericin B (right) in SM versus SM+HBMM. **(F)** Fluconazole IC_50_ values after 24h and corresponding growth curves in SM and SM+HBMM as in (D). Trailing-like growth at later time points was observed upon supplementation with HBMM in presence of fluconazole. **(G)** Disc-diffusion assays with 100 µg fluconazole applied to central disc and imaged after 48 hours of growth in SM growth media alone (left) or supplemented with HBMM (right). Images were cropped in the middle for better visualisation and comparison between conditions.

Next, we quantified antifungal sensitivity using MDAs in SM alone, or supplemented with HBMM. In the case of 5-FC, the metabolite mix increased the IC_50_ by ∼10-fold compared to SM alone (Fig. 2D). While the HBMM had no strong impact on the resistance of the other classes of antifungals (Fig. 2E, F), it supported growth beyond 24 hours in the presence of high concentrations of FLC (supra-MIC growth, Fig. 2F), indicating azole tolerance, a phenotype distinct from resistance [19]. Disc-diffusion assays, an established method to capture azole tolerance [9] confirmed the effect of HBMM on FLC tolerance, as we observed a high fraction of colonies growing within the zone of inhibition after 48 hours (Fig. 2G) upon supplementation with HBMM. Thus, a supplement of human blood metabolites at physiological concentration affects *S. cerevisiae* physiology and metabolism, increases FLC tolerance and reduces 5-FC efficacy.

### Human metabolites affect FLC tolerance and 5-FC resistance

To assess which and how human blood metabolites affected antifungal efficacy, and to exclude that antagonistic and synergistic interactions were masking potential effects when all metabolites were mixed together, we screened individual metabolite-antifungal pairs using MDAs (Fig. 3A). As a primary read-out for resistance, we determined dose-response curves for each condition to estimate the drug concentration to inhibit 50% of growth (IC_50_) after 24 hours. To approximate tolerance, we additionally calculated the obtained biomass (OD_600_) at a “supra-MIC” concentration (80 µg/ml, 40 hours growth). Again, we observed effects of individual or sets metabolites on fungal susceptibility against 5-FC and FLC (Fig. 3B-D), and not for CSP and AmB (Supplementary Fig. 3A, B).

**Figure 3.**
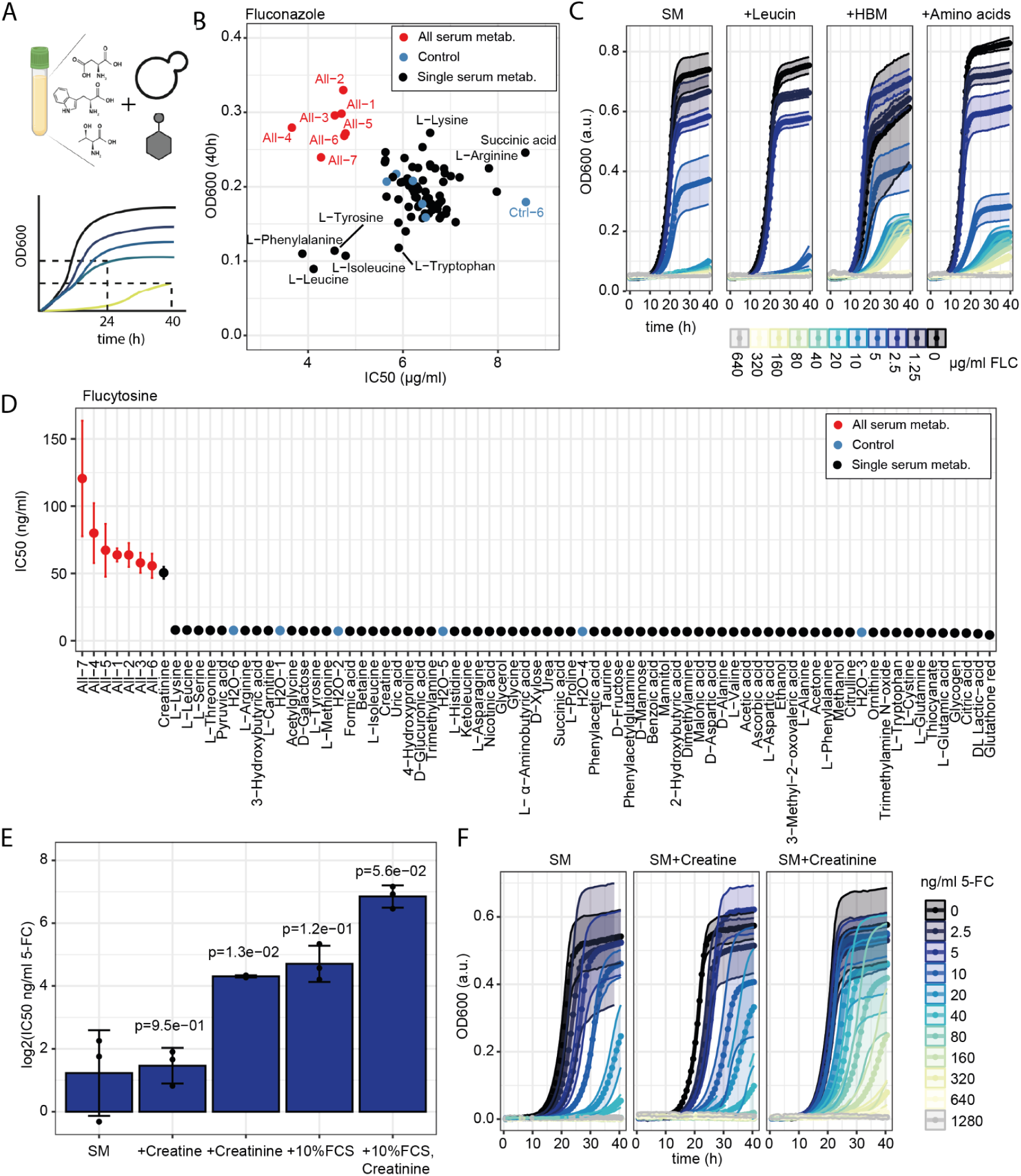
Testing individual antifungal-metabolite interactions reveals creatinine as an antagonist of flucytosine. **(A)** Schematic overview of the antifungal-metabolite interaction screen. Sensitivity of *S. cerevisiae* to an antifungal in SM alone, or in the presence of no (blue), all (red) or individual (black) metabolites at blood-like concentrations, and controls, was assessed by microbroth dilution assays using 6 concentrations of antifungal, and 2 untreated controls, and IC_50_ values were estimated from dose–response curves using OD_600_ after 24h. In addition to overall sensitivity, azole tolerance was estimated using OD_600_ after 40h of growth in the presence of 80 µg/ml fluconazole. **(B)** Screening results as outlined in (A) for the antifungal fluconazole. Overall sensitivity was estimated by IC_50_ values (x-axis) and trailing growth was estimated by OD_600_ after 40 h (y-axis). Individual metabolites such as leucine slightly increased fluconazole efficacy, whereas the combination of the full HBMM (“All”) promoted fluconazole tolerance. **(C)** Validation of screening results. Growth curves of *S. cerevisiae* grown in SM or SM supplemented with the indicated metabolites at physiological human blood concentrations were measured across increasing flucytosine concentrations. Data are shown as the mean of n = 3 replicates +/- SD. **(D)** Screening results as outlined in (A) for the antifungal flucytosine. Shown are IC_50_ values after 24 h of growth, with error bars indicating the lower and upper bounds of the 95% confidence interval derived from the dose–response model. **(E)** *S. cerevisiae* was grown in SM medium with or without the indicated supplements (10% FCS, 79µM creatinine, 79µM creatine, or the combinations), and antifungal susceptibility was assessed by microbroth dilution with increasing antifungal concentrations. IC_50_ values were obtained based on dose-response curves using OD_600_ after 24h of growth. Data are shown as the mean of n = 3 replicates. Statistics based on t-tests comparing each condition to SM alone, with Holm correction for multiple testing. **(F)** Corresponding growth curves for the data shown in (E), comparing the effect of creatinine supplementation on growth in presence of increasing levels of flucytosine to the untreated control. Data are shown as the mean of n = 3 replicates +/- SD.

In the case of FLC, hydrophobic amino acids such as leucine slightly reduced the IC_50_, by up to 2-fold (Fig. 3B, C). This effect was masked by supplementation with the full HBMM, which again induced supra-MIC growth compared to untreated controls. No single compound induced comparable tolerance at physiological concentration. This finding is in line with a previous study, where we found that rich extracellular metabolomes produced by synthetic communities of *S. cerevisiae* lead to azole tolerance by promoting drug export [21]. Notably, we find that these interactions also occur at physiologically relevant conditions, as supplementing all in blood abundant proteinogenic amino acids at physiological concentration similarly induced tolerance (Fig. 3C).

Turning to 5-FC, we confirmed the strong increase in the IC_50_ of *S. cerevisiae* against 5-FC by HBMM addition. The drop-in screen identified creatinine as the compound that was responsible for the effect (Fig. 3D), and addition of creatinine alone significantly increased the IC_50_, whereas its precursor creatine had no effect (Fig. 3D-F). Further, addition of creatinine also reduced the efficacy of 5-FC in a more complex environment, as simulated by addition of 10% FCS (Fig. 3E). Thus, creatinine, an abundant metabolite in human biofluids, antagonizes the antifungal drug 5-FC.

### Creatinine antagonizes 5-FC uptake and efficacy

In humans, creatine is both an endogenously synthesized and a dietary metabolite. Phosphorylated creatine is involved in rapid, anaerobic regeneration of ATP from ADP, to buffer rapid changes in ATP levels, in particular in muscle [37]. Creatine further undergoes a spontaneous reaction to form the cyclic end-product creatinine (Fig. 4A), which is excreted. We noted that the chemical structure of creatinine resembles that of 5-FC (Fig. 4A). Instead, the structurally dissimilar creatine showed no significant interaction with 5-FC (Fig. 3E, F). We thus hypothesized that creatinine could compete for 5-FC uptake through cytosine-purine permeases (Fig. 1C). We started by quantifying 5-FC upon exposing cells to creatinine. Indeed, creatinine in growth media reduced intracellular 5-FC levels in Δfcy1 cells in a concentration dependent manner (Fig. 4B).

**Figure 4.**
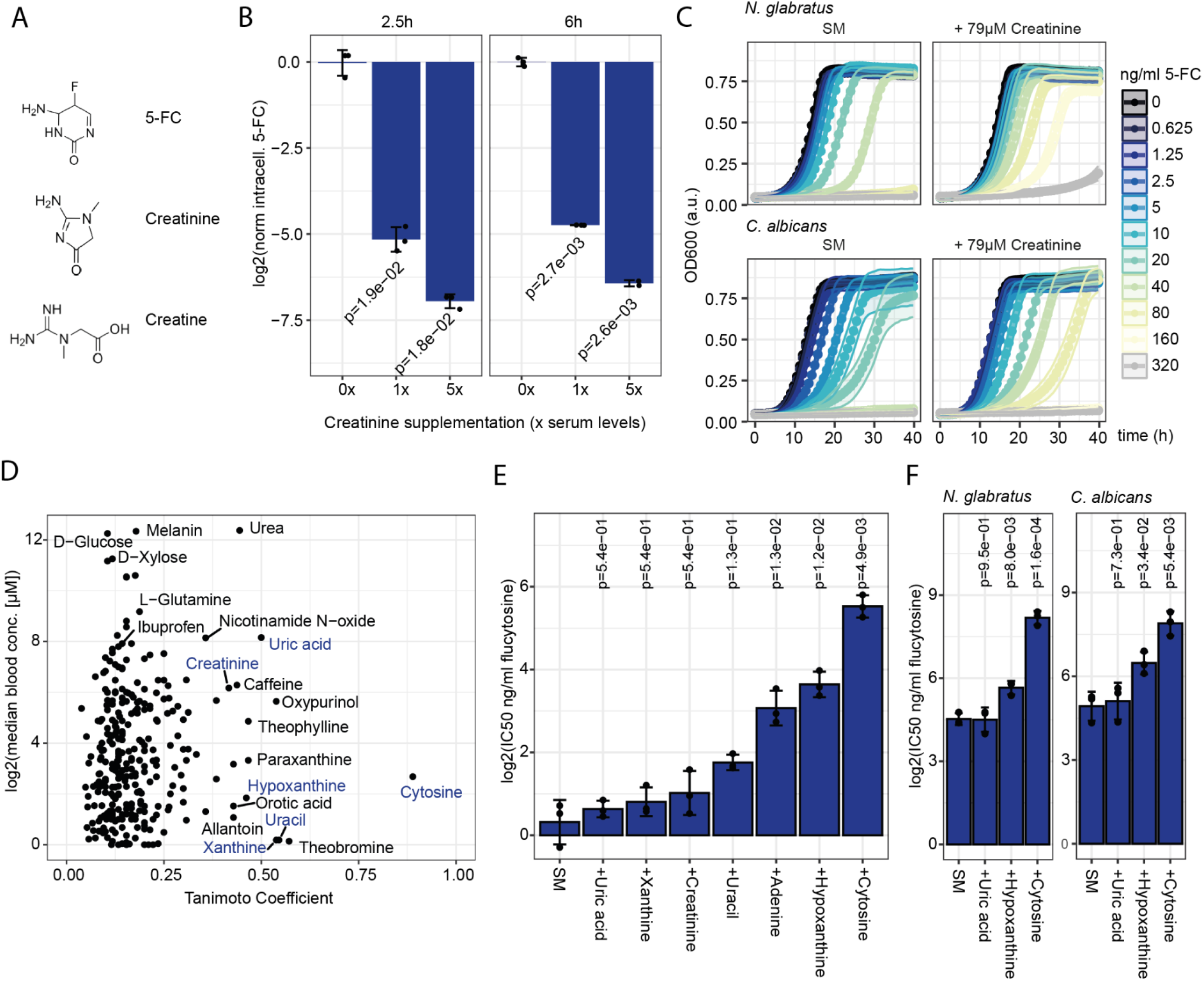
Creatinine impairs flucytosine efficacy by competing for uptake, extending to other host-associated metabolites and yeast pathogens. **(A)** Chemical structures of flucytosine, creatinine, and creatine. **(B)** Supplementation of SM medium with x-fold serum levels of creatinine (79 µM) reduced intracellular flucytosine levels, as estimated by LC-MS/MS, in Δ*fcy1* cells after 2.5 and 6 h of growth in a dose-dependent manner. Statistics were based on t-tests comparing each creatinine condition to the unsupplemented control, with Holm correction for multiple testing. **(C)** Growth curves from microbroth dilution assays with flucytosine for the human fungal pathogens *N. glabratus* and *C. albicans* comparing creatinine supplementation to the untreated control. Data are shown as the mean of n = 3 replicates +/- SD. **(D)** Chemical similarity of the antifungal flucytosine to 292 metabolites with relaxed filters and estimated concentrations >1µM in human blood. Shown are metabolite blood concentrations (y-axis) and the Tanimoto coefficient as a measure of chemical similarity (x-axis). Compounds selected for testing for flucytosine antagonism are highlighted in blue. (E) *S. cerevisiae* was grown in SM medium with or without the indicated metabolites (at 10µM) that share structural similarity with flucytosine, and flucytosine susceptibility was assessed by microbroth dilution with increasing flucytosine concentrations. Corresponding IC50 values were estimated from dose–response curves using OD600 after 24h. Shown are mean values +/- SD of n=3 replicates. Statistics were based on t-tests comparing each condition to SM alone, with Holm correction for multiple testing. **(F)** As in (E), select metabolites were tested for antagonism against flucytosine in *N. glabrata* (left) and *C. albicans* (right).

Next, we tested this antagonism of creatinine on 5-FC in other *S. cerevisiae* isolates, as well as the major clinical yeast pathogens causing invasive infections with high mortality [38], *Candida albicans* and *Nakaseomyces glabratus.* In all sensitive strains, creatinine antagonized the effects of 5-FC (Fig. 4C, Supplementary Fig. 4A). We wondered if other metabolites could have a similar effect, including drugs and lower abundance serum metabolites not included in the HBMM. Comparing estimated median metabolite concentration of HMDB blood-metabolite dataset to the chemical similarity with 5-FC, as quantified by the Tanimoto coefficient [39], revealed several metabolites of structural similarity (Fig. 4D). Cytosine had the highest structural similarity, followed by uracil, and purines including uric acid, adenine, xanthine, hypoxanthine or dietary compounds such as caffeine or drugs like oxypurinol, and creatinine. Testing the effect of selected metabolites in *S. cerevisiae* at a fixed concentration of 10 µM confirmed that cytosine most strongly antagonized 5-FC, followed by hypoxanthine, uracil, and creatinine (Fig. 4E). While chemical similarity with 5-FC explained antagonism for most metabolites, xanthine, as well as uric acid had no effect across tested species (Fig. 4E, F). We speculated that their 2-carbonyl group prevented the interaction with permeases that transport 5-FC, particularly for the major permease Fcy2p involved in 5-FC import (Fig. 1C, D) [29]. Taken together, compounds recognized by purine–pyrimidine permeases most strongly affected the fungistatic activity of 5-FC. Cytosine and adenine, though usually present at low abundance, displayed strong antagonistic effects, requiring ∼10-fold and ∼30-fold stoichiometric excess, respectively, to counteract 5-FC. By contrast, creatinine showed weaker antagonism (Supplementary Fig. 4B). Yet as creatinine is abundant in human biofluids, it likely more strongly inhibits 5-FC uptake under host conditions.

### Creatinine strongly inhibits 5-FC activity against *N. glabratus* in human urine

Creatinine levels differ between biofluids, with extremely high levels in human urine (Fig. 5A), where 5-FC can be used for the treatment of urinary candidiasis caused by azole-resistant *N. glabratus [40,41]*. However treatment can fail and secondary resistances are frequently observed [41]. Curiously, differences in the fungistatic activity and rate of resistance evolution against 5-FC have long been reported between serum and urine [42,43]. We hence suspected that this discrepancy could be attributable to high creatinine levels in urine. To test this hypothesis, we focused on a *N. glabratus* isolate (BG2), which was susceptible to 5-FC in minimal media (IC_50_ ∼0.32 µM). At blood levels of creatinine, the IC_50_ increased to 1.6 µM, and when adding the concentration typical for urine (∼9.8 mM Creatinine) to 52.2 µM, exceeding the suggested 5-FC epidemiological cutoff value of >4 µM for *N. glabratus* [15] (Fig. 5B).

**Figure 5.**
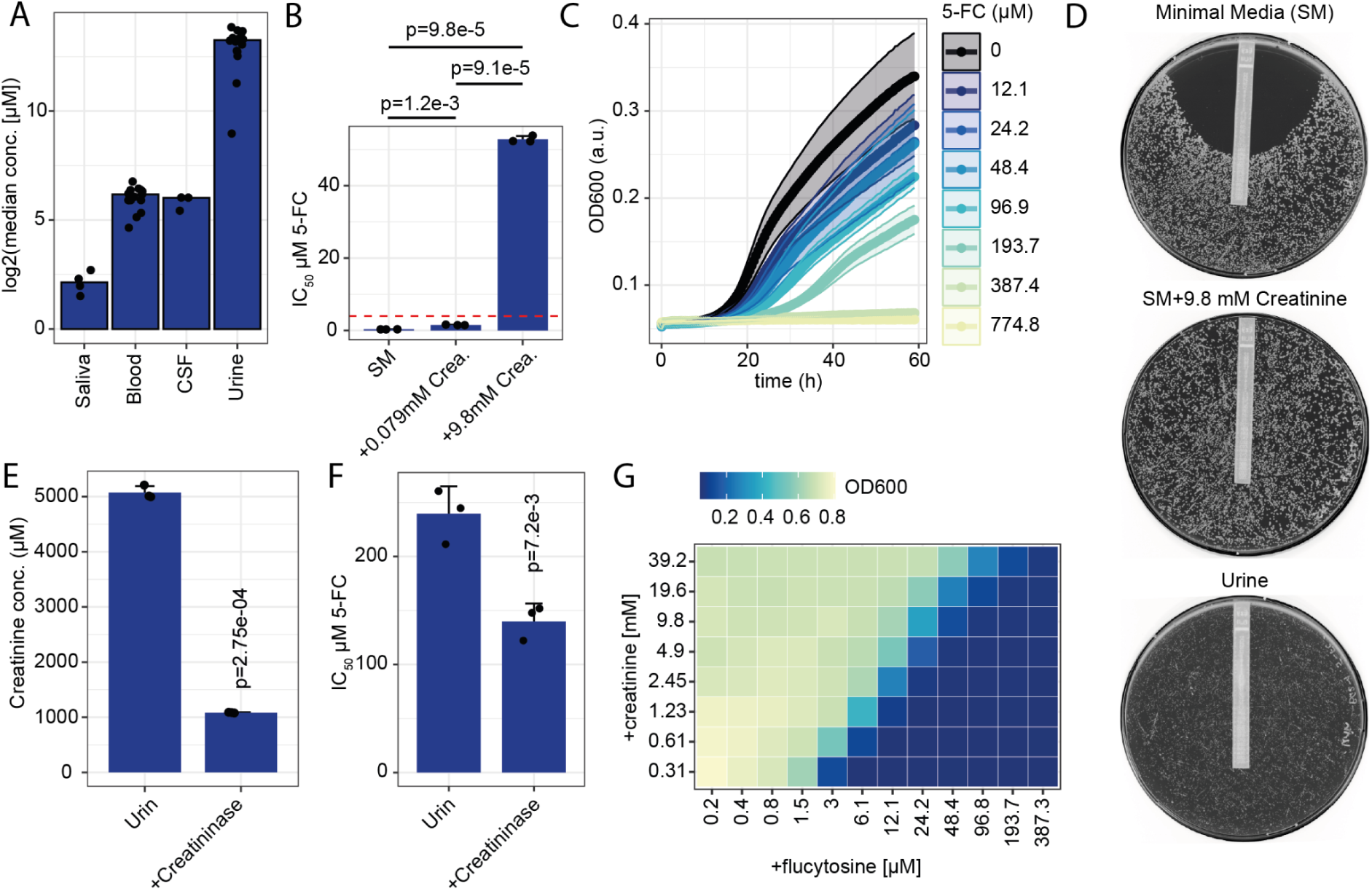
Creatinine is a key component affecting flucytosine efficacy in urine. **(A)** Estimated creatinine concentrations in human biofluids, compiled from the Human Metabolome Database [32]. Each point represents the reported estimated concentration from an individual study, and bars indicate the median creatinine concentration for each biofluid across all studies. **(B)** *N. glabrata* was grown in SM medium alone or supplemented with creatinine at blood (79µM) or urine (9800µM) concentrations. Flucytosine susceptibility was assessed by microbroth dilution across increasing drug concentrations, and IC_50_ values were estimated from dose–response curves using OD_600_ after 24 h. Data are shown as the mean of n = 3 replicates. Statistics were based on t-tests for the indicated comparisons. **(C)** Growth curves of *N. glabratus* grown in human urine with increasing flucytosine concentrations. Note that *N. glabrata* grew poorly in urine, requiring extended incubation times to estimate IC_50_ values. **(D)** Disc diffusion was performed using EStrips containing a flucytosine gradient which was placed on minimal medium alone, minimal medium supplemented with urine-level creatinine, or human urine supplemented with 0.1% glucose. Images were acquired after 6 days of incubation at 30 °C to compensate for slow growth in urine. Representative images from n=2 independent replicates. **(E)** Creatinine levels in untreated urine or urine treated with creatininase after 2 h of incubation at 30 °C. Data are shown as mean +/- SD of n=3 replicates; the indicated p-value was calculated using a t-test. **(F)** *N. glabratus* was grown in urine supplemented with 0.1% glucose and pre-treated with creatininase or the corresponding control as indicated in (E), and flucytosine susceptibility was assessed by microbroth dilution across increasing flucytosine concentrations. IC_50_ values were estimated from dose–response curves using OD_600_ after 30h to compensate for slower growth in urine compared to growth media. Data are shown as mean +/- SD of n = 3 replicates. Statistics were based on t-tests comparing each condition to the corresponding control. **(G)** Heatmap of a checkerboard-like assay showing OD_600_ of *N. glabratus* after 24 h of growth across increasing creatinine (y-axis) and flucytosine (x-axis) concentrations. Data based on the mean of n=3 replicates.

Next, we directly tested human urine (sample from Innovative Research), in which *N. glabratus* poorly grew and required longer incubation times. Urine proved highly antagonistic to 5-FC (Fig. 5C). A disc diffusion assay using a concentration gradient with the Etest method yielded similar results: addition of creatinine or growth in human urine directly rendered the drug ineffective up to the highest concentration of the test in the otherwise susceptible isolate (Fig. 5D). To further confirm that creatinine was antagonizing 5-FC efficacy even at such high levels in urine, we set up an enzyme assay, using a recombinant creatinine amidohydrolyase (creatininase) in urine and measured creatinine by LC-MS/MS. First, we quantified creatinine levels in the urine sample, and detected concentrations around ∼ 8 mM. Adding creatininase quickly reduced creatinine levels, but reached a plateau at ∼ 2 mM (Supplementary Fig. 5A). Presumably this is due to product-inhibition by the formed creatine, as additional creatine or creatinine led to a marked increase in remaining creatinine in urine after creatininase treatment (Supplementary Fig. 5B). Importantly, addition of the antifungal 5-FC had no effect on creatinine levels, and the drug was not degraded by creatininase in urine (Supplementary Fig. 5A). This allowed us to directly assess the impact of the reduction of creatinine on the fungistatic effect of 5-FC. We thus performed a microbroth dilution assay with flucytosine in another urine sample, with or without creatininase-treatment. Reassuringly, in line with reduction of creatinine levels (Fig. 5E), the IC_50_ of 5-FC was reduced in creatininase-treated urine (Fig. 5F). To further validate that creatinine antagonises flucytosine at such high concentrations, we tested the uncovered creatinine-flucytosine interaction in minimal media by ramping the estimated therapeutic range of 5-FC in serum [44] against and above healthy urine creatinine levels. The fungistatic effect of 5-FC decreased linearly, with a doubling of creatinine increasing the IC_50_ by roughly two-fold (Fig. 5G). In summary, creatinine antagonizes 5-FC efficacy in N. glabratus at physiologically concentrations in human urine.

## DISCUSSION

Metabolism and biochemical context is key in understanding host-pathogen interactions, as well as drug responses. While it is broadly accepted that competition for nutrients and metal ions are important during infection, a growing body of evidence suggests that the biochemical environment also influences infection-relevant phenotypes, such as the efficacy of antimicrobials [20–22,24,45]. However, because of the complexity of the metabolomes in biofluids and at different infection sites, we often don’t know which metabolites are causal for the observed phenotypes. Particularly in fungi, knowledge has been sparse and limited to few conditions, compounds, and to alterations with strong physiological impact, such as changing carbon source or pH [23,24], or interactions with other drugs [28].

Here, we show that altering the biochemical environment, even by supplementation with small molecules that do not primarily fuel growth, modulates the efficacy of main antifungals. With an emphasis on 5-FC, we identify an underlying mechanism in the direct interference of a host-derived metabolite, creatinine, with drug uptake. This interaction alters 5-FC efficacy, both in budding yeast, that we used as a model, as well as in *C.albicans* and *N glabratus*. Our results confirm a long postulated competition between canonical substrates of purine-pyrmidine permeases such as hypoxanthine with the drug 5-FC [31], and further show that this competition also involves seemingly unrelated molecules such as creatinine. Of note, as purines are no likely substrate of Fcy1p-mediated desamination, and as hypoxanthine lacks a free amine altogether, our data reinforces the hypothesis that inhibition of uptake is the major mode of action, and not antagonism of the desamination reaction of 5-FC by Fcy1p. Furthermore, we show that these competitions occur at concentrations that are physiologically relevant at infection sites, particularly the urinary tract. Notably, this mechanism may extend to other hydrophilic antimicrobials that rely on specific import systems [46]. The impact of environmental factors on antifungal efficacy primarily affects FLC and 5-FC, two drugs that target intracellular processes; similarly key emerging antifungals also act intracellularly [11]. Instead, we observed less environmental influence on the efficacy of AmB or CSP, which target membrane and cell wall, respectively [11].

5-FC has high in vitro activity and in vivo bioavailability [18]. However, it is rarely used as monotherapy, partly because of secondary resistance and high rates of treatment failure e.g. for urinary candidiasis [41]. Indeed, in vitro studies have found 5-FC to be less efficacious and leading to higher rate of acquired resistance compared to serum [42,43]. According to our results, this discrepancy in efficacy could be due to high creatinine levels in urine, and testing how this would shape the evolutionary trajectory and dynamics of resistance evolution will be important in the future.

Notably, the detected increase in antifungal resistance and tolerance manifest within recommended treatment levels, which for 5-FC range from >20 mg/mLl (>155 µM) to <100 mg/mL (<775 µM), where drug levels are reduced to avoid hepatotoxicity [41,47]. However, 5-FC levels vary substantially between patients, and are often lower [44]. While the concentration in urine should generally exceed serum levels due to creatinine-like clearance [18], we only found data from the 70s, where with then applied treatment regimes 5-FC levels in urine ranged in patients and sampling periods between 0 and 4.2 mg/mL (∼32.5 µM) [43,48]. These observations align with the generally positive outcomes of 5-FC treatment for urinary candidiasis [41], as urinary 5-FC concentrations typically exceed the IC_50_ predicted from creatinine alone. Nevertheless, levels can approach or surpass the observed IC_50_ in urine (∼50 µM), where creatinine and other antagonistic metabolites may reduce drug uptake. Moreover, metabolite concentrations can fluctuate: for example, cytosine increases with certain diets [49], and creatinine can rise due to supplements or altered kidney function [50]. These factors suggest that in individuals with low 5-FC availability, elevated antagonists could compromise efficacy or favor the emergence of secondary resistance, potentially contributing to variability in treatment outcomes.

Thus, as exemplified by the interference of creatinine with flucytosine, host factors can directly modulate antifungal efficacy across fungal species. These findings highlight the need to incorporate relevant host-derived components into in vitro models, particularly in clinical susceptibility testing, to better capture such interactions [22,45]. More broadly, the approach employed here–reconstituting human biofluids in vitro based on knowledge of their molecular composition–is increasingly recognized for its potential to enhance physiological relevance, yet remains relatively underused. Particularly in microbiology, such approaches enable scalable phenotyping in environments that better approximate those encountered during infection, these models may improve the prediction of in vivo drug efficacy, narrowing the gap between in vitro laboratory susceptibility testing and clinical outcomes.

## ACKNOWLEDGEMENTS

We thank Daniela Ludwig, Christiane Kilian, Lukasz Szryviel (all Charité - Universitätsmedizin Berlin, Berlin), as well as Michael Mülleder (High-Throughput Mass Spectrometry, Charité - Universitätsmedizin Berlin). We further acknowledge Lynn Jade Nai Chung Tong (University of Applied Sciences, Leiden) and Artem Khan (Rockefeller University, New York) for support and discussions in early stages of this project. We also thank Austin Mottola and Julia Muenzner (both Charité - Universitätsmedizin Berlin) and Judy Berman (Tel Aviv University) for critical and helpful discussion of the manuscript. We further thank Judy Berman for generously gifted yeast strains. This work was supported by the Swiss National Science Foundation Postdoc.Mobility fellowship 191052 (to J.H), and the European Research Council with an ERC-SyG-2020 951475 (to M.R).

## DATA AVAILABILITY

Data that support the findings of this study are available within the article or the supporting information. Any additional raw data are available without restriction upon request from the corresponding authors.

## METHODS

### Cultivation, growth media, and components

All growth experiments were performed at 30 °C unless otherwise indicated. Synthetic Minimal (SM) medium consisted of 6.7 g/L Yeast Nitrogen Base (YNB) with ammonium sulfate and without amino acids (Becton Dickinson), and 5 g/L glucose (Sigma-Aldrich). Synthetic Complete (SC) medium was prepared by supplementing SM medium with 0.56 g/L Complete Supplement Mixture (CSM; mbio). RPMI medium was prepared using 4.12 g/L RPMI-1640 without L-glutamine and sodium bicarbonate powder and 5 g/L glucose (both Sigma-Aldrich). YPD medium was composed of 20 g/L peptone, 10 g/L yeast extract (both Becton Dickinson), and 5 g/L glucose (Sigma-Adlrich). All media were buffered with 0.1 M MOPS (Sigma-Aldrich) adjusted to pH 6.75. All media were sterilized by filter sterilization. Where indicated fetal calf serum (Biochrome) or pools of normal human serum or human urine (from Zen-Bio, Innovative Research) were sterile filtered and added to SM medium or used directly. For solid cultures, an autoclaved, 50g/L agar solution was diluted with respective media to a final concentration of 20 g/L agar (Becton Dickinson). Antifungal compounds were obtained from Sigma-Aldrich unless otherwise noted. Fluconazole, caspofungin, amphotericin B, and flucytosine were dissolved in DMSO and stored at –20 °C. From these stocks, solutions with working concentrations were freshly prepared prior to each experiment.

### Yeast strains

Unless otherwise stated, experiments were performed with the haploid *S. cerevisiae* strain BY4741ki, in which the common auxotrophies (*his3*, *leu2*, *met15*, *ura3*) have been complemented with genomic knockin of *HIS3*, *LEU2*, *URA3* and *MET15* with native promoters [21]. Additional *S. cerevisiae* strains used in this study are from the haploid deletion library where protrophy was restored with the plasmid pHLUM [51], including BY4741 Δ*fcy1*, Δ*fcy2* and Δ*fcy21*. The corresponding deletion strain BY4741 Δ*his3* was used as a control strain. Additional isolates used include the *S. cerevisiae* strains BCC, BDT, BME, BVD, CIR, CKR and YAN from the 1011 *S. cerevisiae* isolates collection [52]. Other yeast species used in this study were *C. albicans* strain SC5314, and *N. glabratus* strain BG2 (both gifted from Judy Berman, Tel Aviv University).

### Growth curves and microbroth dilution assays (MDAs)

Strains stored as glycerol stocks were streaked on YPD agar plates and incubated at 30 °C up to 3 days. Individual colonies were used to inoculate 10 mL of the respective liquid media in Erlenmeyer shake flasks and grown overnight at 30 °C with shaking at 300 rpm. Subsequently, the overnight cultures were diluted ∼1:20 in fresh media and incubated for an additional 4 hours. The optical density at 600 nm was then adjusted to 0.001. For drug sensitivity assays, the cell suspension was mixed in a 1:1 v/v ratio to create a 2-fold serial dilution series of the respective antifungal in a 96-well microtiter plate. To increase the sensitivity of the assay, for the MDA in human urine treated with creatininase, a 1.33-fold serial dilution was used. Each plate included a no-drug and media-only control (blank). To minimize evaporation Nunc Edge (Thermo Fisher Scientific) plates were used, filling the outer rims with a total of 6 mL.

Growth dynamics were assessed by repeated measurements of absorbance at 600 nm over up to 3 days at 30 °C. Measurements were taken with an Infinite 200 PRO® automated plate reader (Tecan) operating in “smooth” mode. Multi-well reads were performed using a 3×3 pattern with a border distance of 1500 µm, and mean values were used for subsequent analysis. Alternatively, growth data were acquired using a Spark-Stack™ (Tecan) plate reader housed in a 30 °C warm room and operated in kinetic mode for sequential acquisition of multiple 96- or 384-well plates. This instrument was configured for multi-well reads using a 3×3 pattern with a border distance of 2000 µm, with mean values used for analysis. For 384-well plates, single-well reads were acquired. Unless otherwise indicated, IC_50_ was based on OD_600_ values after 24h of growth. Dose response curves were fitted using the LL.3 function unless otherwise stated from the R package *drm* using non-normalised OD_600_ values after 24 hours of growth including controls without, and a dilution series of respective antifungal concentrations. IC_50_ values, i.e. the concentration at which 50% of growth was inhibited, were estimated using the R package *ec50estimator*. Growth rates were estimated using the R package grofit [53] based on the log2 fold-change of the ratio between the OD_600_ at each time point and the initial OD_600_ measurement.

### High-throughput microbroth dilution assays to test metabolite-antifungal interactions

Exponentially growing *S. cerevisiae* BY4741ki cultures were prepared in synthetic medium (SM). Then, drug dilutions were prepared in 1.25-fold concentrated SM (8.375 g/L yeast nitrogen base with ammonium sulfate, 0.625% w/v glucose, and 125 mM MOPS) containing cells at an OD_600_ of 0.00125. In addition to control plates without antifungals, the following concentration ranges were tested: fluconazole (80, 40, 20, 10, 5, 2.5 µg/mL), flucytosine (160, 80, 40, 20, 10, 5 ng/mL), caspofungin (8, 4, 2, 1, 0.5, 0.25 µg/mL), and amphotericin B (2.5, 1.25, 0.625, 0.3125, 0.156, 0.078 µg/mL). In parallel, serum-metabolite plates were prepared in 200 µL 96-well microtiter plates, with each well containing either an individual serum metabolite, a water control, or a mixture of all serum metabolites, each at 5-fold physiological concentration (see Supplementary Table 1). A volume of 40 µL from each well was transferred into new 96-well plates using a Beckman i7 96-head pipetting robot. Subsequently, 160 µL of the above-described 1.25-fold medium–drug–cell suspension was added to each well and each plate by robotic pipetting, mixed briefly, and dispensed in order from no-drug to highest drug concentration to minimize tip usage. Each 96-well plate corresponded to one drug concentration, including two no-drug controls per antifungal. From sets of four 96-well plates, 75 µL per well was transferred into 384-well plates using the pipetting robot, again in order from no-drug to low-drug to high-drug concentrations. For each drug, this resulted in two 384-well plates, with each plate containing three drug concentrations and a no-drug control. Growth data were acquired on a Spark-Stack™ (Tecan) plate reader using continuous single-well OD_600_ measurements and analyzed as described above for regular MDAs. For technical replicates of “no drug” controls, means were calculated prior to analysis, and data from L-cysteine was removed as it was highly variable between replicates.

### Metabolite concentrations in human body fluids

The data package serum_metabolites.xml was obtained from human metabolome database [32] (hmdb.ca, access, 10.09.2020) and filtered using the following settings: indicated biospecimen had to be blood, subjects had to be adult and healthy, metabolites had to be quantified in at least n=2 studies. If a concentration range was given, the conservative lower abundance estimate was used. Further, lipids and anorganic molecules were removed from analysis. Only abundant metabolites with a median blood concentration across studies above 10 µM were acquired for testing. Resulting data was further manually curated to remove drugs, insoluble compounds, metabolites we assumed were primarily intracellular such as ATP, metabolites that were not commercially available, and metabolites that were already contained in SM medium, yielding a total of 67 metabolites. All compounds were prepared as concentrated stock solutions (see Supplementary Table 1) and stored at -20 °C, except for L-Cysteine, Glutathione, and Urea which were freshly prepared prior to experiments. To estimate creatinine concentration in cerebrospinal fluid, blood, saliva, and urine, the respective body fluids were searched via hmdb.ca and filtered for data from healthy adults only.

### Chemical similarity analysis

Metabolites were acquired from hmdb.ca as described above, but with relaxed filters, where blood metabolites with a median concentration above 1 µM from healthy adults were considered even if they were only detected in n=1 study. Besides removing lipids and inorganic molecules, no additional filters or manual curation was applied, yielding a total of 292 metabolites. Using the fmcsBatch function of the fmcsR package, the similarity of each of these metabolites to flucytosine (5-FC) was quantified by calculating the corresponding Tanimoto Coefficient using the maximum common substructure (MCS) algorithm [54].

### Disc diffusion and Etest assay

Strains stored as glycerol stocks were streaked onto YPD agar plates and incubated at 30 °C for 2–3 days. Single colonies were used to inoculate 10 mL of the respective liquid media in Erlenmeyer flasks, which were then grown overnight at 30 °C with shaking at 300 rpm, and were then diluted in fresh media and incubated for additional 4-6 h. Exponentially growing cells were then diluted in water to an OD_600_ of 0.005 and 100 µL of cell suspension was distributed on 9 cm petri dishes filled with 20 mL solid media using 4 mm glass beads, which were subsequently removed. Then, a 6 mm disc was placed in the center of the plate and 10 µL of 10 µg/mL fluconazole were added. Following incubation at 30 °C plate imaging was performed at the indicated timepoints, typically after 24h, 48h and 72h using Epson Perfection V800 Photo (Epson) scanner using a 2×2 stencil and following default settings using a previously described custom made software (*pyphe)* [55]. Colors of images were subsequently inverted for better visualisation. For the Etest method, the experiment was performed in the same fashion, but a commercial Ezy MIC™ Strips (HIMEDIA) containing an antifungal gradient was placed in the center of the agar plate containing cells.

### Sample preparation for intracellular flucytosine quantification

Exponentially growing *S. cerevisiae* strains as indicated were inoculated in 40 mL SM containing 0.64 µg/mL flucytosine at a starting OD_600_ of 0.008. Cultures were incubated shaking at 30 °C and were harvested. At each indicated time point, a volume of culture corresponding to an OD_600_ of 2 was pelleted by centrifugation, the majority of the supernatant was carefully removed, and the pellet was resuspended in ∼50 µL of residual supernatant. In order to rapidly remove extracellular flucytosine, a fast-filtration setup was adapted [56]. Briefly, a 47 mm radius, 0.45 µm size-cutoff nylon membrane filter (Merck Millipore) was pre-conditioned with 30 mL MilliQ and was mounted on a vacuum filtration funnel connected to a vacuum flask. Cell suspensions were pipetted onto the filter, washed with 100 mL Milli-Q water, and the filter was immediately transferred into 5 mL tubes prefilled with 600 µL MS grade water containing 1 µM stable isotope labelled 5-FC as internal standard. Filters were vortexed vigorously for ∼30 s, and 400 µL of the suspension was transferred into 2 mL tubes preloaded with 1200 µL 75% ethanol for extraction. Tubes were incubated at 80 °C for 6 min with intermediate vortexing steps. Samples were then centrifuged (12000 x g, 3 min) to pellet debris, and the supernatants were collected into fresh tubes. Solvent was removed in a SpeedVac concentrator, pellets were resuspended in 200 µL 75% ethanol shaking at 1200 rpm on an Eppendorf mixer for 10 min. After centrifugation (12,000 x g, 3 min), 150 µL of the supernatant was used for LC-MS/MS analysis.

### Creatinine quantification in urine and creatininase assays

Frozen human urine samples were obtained from Innovative Research. Where indicated, flucytosine was added from a concentrated stock to a final concentration of 150 µM to assess its stability in the presence of creatininase. Similarly, where indicated, additional creatine or creatinine was spiked into urine samples to assess their effect on creatininase activity. Creatininase (from *Pseudomonas putida*; MedChemExpress) was prepared as an approximately 20x concentrated stock in water, and 5, 10, or 20 units were added to 40 µL urine, or to urine supplemented with creatine, creatinine, or flucytosine as indicated. Samples were collected over time; the first timepoint was taken on ice, after which samples were transferred to and incubated at 37 °C. Urine samples intended for subsequent microbroth dilution assays were incubated with creatininase (5 units per 40 µL urine) or without enzyme for 2 hours at 37 °C, then frozen until use. For LC-MS/MS quantification, samples were diluted 1:200 in MS-grade water prior to analysis.

### Quantification of creatinine and flucytosine by LC-MS/MS

Compounds were analyzed and quantified by LC-MS/MS using an Agilent 1290 liquid chromatography, hyphenated to a Agilent 6470 triple quadrupole mass spectrometer, following a method adapted from a previously described protocol for various antifungal agents [57]. To this end, 1 µL sample was injected and analyzed by reverse-phase chromatography using an Agilent EclipsePlus C18 RRHD column (2.1×50 mm, 1.8 µm particle size). The mobile phase consisted of 10 mM ammonium formate with 0.1% formic acid in water (phase A) and 0.1% formic acid in acetonitrile (phase B). A multi-step gradient was applied: 1% B (0–0.5 min), increasing to 40% B (0.5–0.7 min), then to 45% B by 2 min; phase B was further increased to 95% over 2-4.9 min, maintained at 95% B from 4.9-5.8 min, returned to starting conditions within 0.1 min and kept until 7.2 min for column equilibration. The column compartment was maintained at 25 °C, with a flow rate of 0.3 mL/min. The mass spectrometer was operating in positive ionization multiple reaction monitoring (MRM) mode.

For creatinine, the transitions 114.06 to 44.1 m/z (CE 21, Fragmentor 90) were used as a quantifier, and 114.06 to 86 m/z (CE 9, Fragmentor 90) as a qualifier. For unlabelled flucytosine, the transition 130.03 to 58 m/z (CE 40, Fragmentor 118) was used as a quantifier and 130.03 to 113 m/z (CE 20, Fragmentor 118) was the qualifier. For isotope labelled flucytosine (2H,15N, Shimadzu, M1097) serving as an internal standard, the transitions 132.03 to 59 m/z, 132.03 to 114 m/z were used, with the same collision settings as used for unlabelled flucytosine. Compounds were identified by matching retention time and transitions obtained using an analytical standard. The instrument was operating with the following source settings: capillary voltage: 3500 V, nozzle voltage 1000 V, gas temp 230 °C, sheath gas temp 300 °C, gas flow 6 l/min, sheath gas flow 8 l/min, nebulizer 50 psi. Data were analyzed by obtaining peak areas by integration using MassHunter Quantitative Analysis (Version B.07.01).

Flucytosine was quantified with spike ins of 500 nM of stable isotope labelled standard and by calculating the ratio of areas of unlabelled over stable-isotope labelled flucytosine as input. Where indicated, absolute quantification was performed using unlabelled flucytosine with a concentration range of 0.25 nM to 5 µM with a linear fit and using the isotope ratio to 500 nM labelled flucytosine as input. To estimate intracellular quantities, it was assumed that 1 mL of culture with OD_600_ of 1 translates to 3.2*10^7 *S. cerevisiae* cells, and that one cell has an intracellular volume of 45.5 fL [58]. Creatinine quantification was performed via external calibration with unlabelled analytical standards measured in technical duplicate ranging from 1.56 to 100 µM as obtained using a 1:1 v/v serial dilution. Calibration curves employed a blank offset and were based on a power curve fit.

### LC MS/MS-based proteomics and data analysis

A cryostock of BY4741ki was streaked onto agar plates. Replicate colonies were inoculated into 200 µL of the respective culture medium in 96-well plates and incubated overnight at 30 °C without shaking. Subsequently, 160 µL of each preculture was transferred to 2-mL wells of a 96-deep-well plate containing 1440 µL of minimal medium and one 2-mm borosilicate bead per well. Plates were sealed with a Breathe-Easier sealing membrane (Sigma Aldrich) and incubated on four orbital shakers (Heidolph Titrama× 1000) at 750 rpm, 30 °C for 8 h. 1.4 mL of culture from each well was harvested by centrifugation (5 min, 4000 x g). Supernatants were discarded, plates were sealed with adhesive aluminum foil, and cell pellets were stored at −80 °C until further processing. Frozen pellets were then processed as described previously [59]. Briefly, to each well, acid-washed glass beads (∼100 mg per well) were added using a custom pre-filled plate dispenser, followed by centrifugation (0.5 min, 4 °C, 1000 x g). Cells were resuspended in 200 µL freshly prepared 7 M urea with 0.1 M ammonium bicarbonate (ABC) per well. Plates were sealed with Cap Mats and lysed via bead milling using a Genogrinder (MiniG, SPEX) for 5 min at 1500 rpm, followed by brief centrifugation (1 min, 4 °C, 3000 x g).

Subsequent processing was performed on semi-automated setup on a Biomek i7 pipetting robot: 20 µL of 5 mM dithiothreitol (DTT) was added to each well, mixed, centrifuged briefly, and incubated for 1 h at 30 °C. After 15 min at room temperature, 20 µL of 5 mM iodoacetamide (IAA) was added, mixed, centrifuged briefly, and incubated in the dark for 30 min at room temperature. Reduced and alkylated samples were diluted with 1 mL 0.1 M ABC, mixed, and centrifuged briefly. Then, 500 µL of the diluted lysate was transferred to a plate containing 2 µg trypsin/LysC per well and incubated overnight at 37 °C. The tryptic digest was stopped by addition of 25 µL of 20% formic acid, and peptides were purified using solid-phase extraction in 96-well format. For clean-up, plates were conditioned with 200 µL methanol, washed twice with 200 µL 50% acetonitrile/water, and equilibrated twice with 0.1% formic acid in water. Samples (500 µL per well) were loaded and washed four times with 200 µL 0.1% formic acid, followed by a final centrifugation step at 250 x g. Peptides were eluted in three steps of 110 µL 50% acetonitrile/water and dried to completeness using a vacuum concentrator. Dried peptides were redissolved in 40 µL 0.1% formic acid.

Samples were analyzed on a Bruker timsTOF Pro mass spectrometer, coupled to a Dionex Ultimate 3000 µsystem (Thermo Fisher Scientific). Prior to LC-MS analysis, 1 µg peptides were chromatographically separated with a 30 min gradient on a Waters HSS T3 column (300um x 150mm, 1.8um) heated to 40 °C, using a flow rate of 5 µL/min where mobile phase A & B are 0.1% formic acid in water and 0.1% formic acid in acetonitrile, respectively. The active gradient increases from 2% to 40% B in 30min. For diaPASEF acquisition, the electrospray source (Bruker Apollo II source, Bruker Daltonics) was operated at 4500 V of capillary voltage, 5.0 l/ min of drying gas and 200 C° drying temperature The dia-PASEF windows scheme was as follows: ion mobility range from 1/K0 = 0.6 to 1.60 Vs/cm2 using equal ion accumulation and ramp times in the dual TIMS analyzer of 100 ms, with each cycle time being 0.5 s. The collision energy was lowered as a function of increasing ion mobility from 59 eV at 1/K0 = 1.6 Vs/cm2 to 20 eV at 1/K0 = 0.6 Vs/cm2. For all experiments, TIMS elution voltages were calibrated linearly to obtain the reduced ion mobility coefficients (1/K0) using three Agilent ESI-L Tuning Mix ions (m/z, 1/K0: 622.0289, 0.9848 Vs/cm2; 922.0097, 1.1895 Vs/cm2; and 1221.9906, 1.3820 Vs/cm2).

Data were processed in DIA-NN (v2.2) [60] using default settings except that the quantification strategy was set to “legacy”; peptides/proteins were identified in library-free mode using the UniProt reference proteome UP000002311 (*S. cerevisiae* S288c). The unique genes matrix output with summarized estimated protein abundances was used for downstream statistical analysis. Differential expression analysis was performed in R. Proteins with >50% missing values across conditions were removed prior to testing. Pairwise comparisons between media (SM, SM+HBMM) were assessed using (two-sided) t-tests on log2 transformed protein abundance values, and p-values were adjusted for multiple testing using the Benjamini–Hochberg false discovery rate (FDR); features with BH-adjusted p-value ≤ 0.05 were considered significant. For pathway enrichment, significantly changing proteins were tested for over-representation against a custom background defined as all quantified proteins with adjusted p-values, using the gprofiler2 gost R package with FDR correction (threshold 0.05) and organism set to *S. cerevisiae*. KEGG and TF terms were retained for visualization, and redundant terms were removed manually prior to plotting.

## Notes

### Competing Interest Statement

M. Ralser is founder and shareholder of Eliptica Ltd. The other authors declare no competing interests.

## REFERENCES

1. Brown GD, Denning DW, Gow NAR, Levitz SM, Netea MG, White TC. Hidden killers: human fungal infections. Sci Transl Med. 2012;4: 165rv13.

2. Salmanton-García J, Cornely OA, Stemler J, Barać A, Steinmann J, Siváková A, et al. Attributable mortality of candidemia - Results from the ECMM Candida III multinational European Observational Cohort Study. J Infect. 2024;89: 106229.

3. Denning DW. Global incidence and mortality of severe fungal disease. Lancet Infect Dis. 2024. doi:10.1016/S1473-3099(23)00692-8

4. WHO. WHO fungal priority pathogens list to guide research, development and public health action. 2022. Licence: CC BY-NC-SA 3.0 IGO. Geneva: World Health Organization; 2022. Available: https://www.who.int/publications/i/item/9789240060241

5. Brown GD, Ballou ER, Bates S, Bignell EM, Borman AM, Brand AC, et al. The pathobiology of human fungal infections. Nat Rev Microbiol. 2024. doi:10.1038/s41579-024-01062-w

6. Bongomin F, Gago S, Oladele RO, Denning DW. Global and multi-national prevalence of fungal diseases-estimate precision. J Fungi (Basel). 2017;3. doi:10.3390/jof3040057

7. Gow NAR, Netea MG. Medical mycology and fungal immunology: new research perspectives addressing a major world health challenge. Philos Trans R Soc Lond B Biol Sci. 2016;371: 20150462.

8. Robbins N, Wright GD, Cowen LE. Antifungal Drugs: The Current Armamentarium and Development of New Agents. Microbiol Spectr. 2016;4. doi:10.1128/microbiolspec.FUNK-0002-2016

9. Berman J, Krysan DJ. Drug resistance and tolerance in fungi. Nat Rev Microbiol. 2020;18: 319–331.

10. Kachroo AH, Laurent JM, Yellman CM, Meyer AG, Wilke CO, Marcotte EM. Evolution. Systematic humanization of yeast genes reveals conserved functions and genetic modularity. Science. 2015;348: 921–925.

11. Puumala E, Fallah S, Robbins N, Cowen LE. Advancements and challenges in antifungal therapeutic development. Clin Microbiol Rev. 2024;37: e0014223.

12. Pfaller MA, Diekema DJ, Turnidge JD, Castanheira M, Jones RN. Twenty years of the SENTRY antifungal surveillance program: Results for Candida species from 1997-2016. Open Forum Infect Dis. 2019;6: S79–S94.

13. Schroeder M, Weber T, Denker T, Winterland S, Wichmann D, Rohde H, et al. Epidemiology, clinical characteristics, and outcome of candidemia in critically ill patients in Germany: a single-center retrospective 10-year analysis. Ann Intensive Care. 2020;10: 142.

14. Fisher MC, Hawkins NJ, Sanglard D, Gurr SJ. Worldwide emergence of resistance to antifungal drugs challenges human health and food security. Science. 2018;360: 739–742.

15. Pfaller MA, Diekema DJ. Progress in antifungal susceptibility testing of Candida spp. by use of Clinical and Laboratory Standards Institute broth microdilution methods, 2010 to 2012. J Clin Microbiol. 2012;50: 2846–2856.

16. Doern GV, Brecher SM. The clinical predictive value (or lack thereof) of the results of *in vitro* antimicrobial susceptibility tests. J Clin Microbiol. 2011;49: S11–S14.

17. Zhanel GG, Saunders DG, Hoban DJ, Karlowsky JA. Influence of human serum on antifungal pharmacodynamics with Candida albicans. Antimicrob Agents Chemother. 2001;45: 2018–2022.

18. Felton T, Troke PF, Hope WW. Tissue penetration of antifungal agents. Clin Microbiol Rev. 2014;27: 68–88.

19. Rosenberg A, Ene IV, Bibi M, Zakin S, Segal ES, Ziv N, et al. Antifungal tolerance is a subpopulation effect distinct from resistance and is associated with persistent candidemia. Nat Commun. 2018;9: 2470.

20. Chen L, Tian X, Zhang L, Wang W, Hu P, Ma Z, et al. Brain glucose induces tolerance of Cryptococcus neoformans to amphotericin B during meningitis. Nature Microbiology. 2024;9: 346–358.

21. Yu JSL, Correia-Melo C, Zorrilla F, Herrera-Dominguez L, Wu MY, Hartl J, et al. Microbial communities form rich extracellular metabolomes that foster metabolic interactions and promote drug tolerance. Nat Microbiol. 2022;7: 542–555.

22. Ersoy SC, Heithoff DM, Barnes L 5th, Tripp GK, House JK, Marth JD, et al. Correcting a Fundamental Flaw in the Paradigm for Antimicrobial Susceptibility Testing. EBioMedicine. 2017;20: 173–181.

23. Marr KA, Rustad TR, Rex JH, White TC. The trailing end point phenotype in antifungal susceptibility testing is pH dependent. Antimicrob Agents Chemother. 1999;43: 1383–1386.

24. Ene IV, Adya AK, Wehmeier S, Brand AC, MacCallum DM, Gow NAR, et al. Host carbon sources modulate cell wall architecture, drug resistance and virulence in a fungal pathogen. Cell Microbiol. 2012;14: 1319–1335.

25. Suchodolski J, Krasowska A. Fructose induces fluconazole resistance in candida albicans through activation of Mdr1 and Cdr1 transporters. Int J Mol Sci. 2021;22: 2127.

26. Ballou ER, Avelar GM, Childers DS, Mackie J, Bain JM, Wagener J, et al. Lactate signalling regulates fungal β-glucan masking and immune evasion. Nat Microbiol. 2016;2: 16238.

27. Williams RB, Lorenz MC. Multiple Alternative Carbon Pathways Combine To Promote Candida albicans Stress Resistance, Immune Interactions, and Virulence. MBio. 2020;11. doi:10.1128/mBio.03070-19

28. Obermeier M, Esparza-Mora MA, Heese O, Cohen N, Varma SJ, Tober-Lau P, et al. Non-antifungal medications administered during fungal infections drive drug tolerance and resistance in Candida albicans. J Med Microbiol. 2025;74. doi:10.1099/jmm.0.002046

29. Paluszynski JP, Klassen R, Rohe M, Meinhardt F. Various cytosine/adenine permease homologues are involved in the toxicity of 5-fluorocytosine in Saccharomyces cerevisiae. Yeast. 2006;23: 707–715.

30. Paluszynski JP, Klassen R, Meinhardt F. Genetic prerequisites for additive or synergistic actions of 5-fluorocytosine and fluconazole in baker’s yeast. Microbiology. 2008;154: 3154–3164.

31. Polak A, Grenson M. Evidence for a common transport system for cytosine, adenine and hypoxanthine in Saccharomyces cerevisiae and Candida albicans. Eur J Biochem. 1973;32: 276–282.

32. Wishart DS, Feunang YD, Marcu A, Guo AC, Liang K, Vázquez-Fresno R, et al. HMDB 4.0: the human metabolome database for 2018. Nucleic Acids Res. 2018;46: D608–D617.

33. Psychogios N, Hau DD, Peng J, Guo AC, Mandal R, Bouatra S, et al. The human serum metabolome. PLoS One. 2011;6: e16957.

34. Geyer PE, Kulak NA, Pichler G, Holdt LM, Teupser D, Mann M. Plasma Proteome Profiling to Assess Human Health and Disease. Cell Syst. 2016;2: 185–195.

35. Cantor JR, Abu-Remaileh M, Kanarek N, Freinkman E, Gao X, Louissaint A Jr, et al. Physiologic Medium Rewires Cellular Metabolism and Reveals Uric Acid as an Endogenous Inhibitor of UMP Synthase. Cell. 2017;169: 258–272.e17.

36. Vande Voorde J, Ackermann T, Pfetzer N, Sumpton D, Mackay G, Kalna G, et al. Improving the metabolic fidelity of cancer models with a physiological cell culture medium. Sci Adv. 2019;5: eaau7314.

37. Wyss M, Kaddurah-Daouk R. Creatine and creatinine metabolism. Physiol Rev. 2000;80: 1107–1213.

38. Pfaller MA, Diekema DJ. Epidemiology of invasive candidiasis: a persistent public health problem. Clin Microbiol Rev. 2007;20: 133–163.

39. Bajusz D, Rácz A, Héberger K. Why is Tanimoto index an appropriate choice for fingerprint-based similarity calculations? J Cheminform. 2015;7: 20.

40. Pappas PG, Kauffman CA, Andes DR, Clancy CJ, Marr KA, Ostrosky-Zeichner L, et al. Clinical Practice Guideline for the Management of Candidiasis: 2016 Update by the Infectious Diseases Society of America. Clin Infect Dis. 2016;62: e1–50.

41. Sigera LSM, Denning DW. Flucytosine and its clinical usage. Ther Adv Infect Dis. 2023;10: 20499361231161387.

42. Davies RR, Savage MA. Observations on 5-fluorocytosine and Candida albicans. Sabouraudia. 1974;12: 302–308.

43. Davies RR, Reeves DS. 5-fluorocytosine and urinary candidiasis. BMJ. 1971;1: 577–579.

44. Pasqualotto AC, Howard SJ, Moore CB, Denning DW. Flucytosine therapeutic monitoring: 15 years experience from the UK. J Antimicrob Chemother. 2007;59: 791–793.

45. Sollier J, Basler M, Broz P, Dittrich PS, Drescher K, Egli A, et al. Revitalizing antibiotic discovery and development through in vitro modelling of in-patient conditions. Nat Microbiol. 2024;9: 1–3.

46. Goldman ID. Membrane transport of chemotherapeutics and drug resistance: Beyond the ABC family of exporters to the role of carrier-mediated processes. Clin Cancer Res. 2002;8: 4–6.

47. Stott KE, Hope WW. Therapeutic drug monitoring for invasive mould infections and disease: pharmacokinetic and pharmacodynamic considerations. J Antimicrob Chemother. 2017;72: i12–i18.

48. Schönebeck J, Polak A, Fernex M, Scholer HJ. Pharmacokinetic studies on the oral antimycotic agent 5-fluorocytosine in individuals with normal and impaired kidney function. Chemotherapy. 1973;18: 321–336.

49. Bar N, Korem T, Weissbrod O, Zeevi D, Rothschild D, Leviatan S, et al. A reference map of potential determinants for the human serum metabolome. Nature. 2020;588: 135–140.

50. Williamson L, New D. How the use of creatine supplements can elevate serum creatinine in the absence of underlying kidney pathology. BMJ Case Rep. 2014;2014. doi:10.1136/bcr-2014-204754

51. Mülleder M, Capuano F, Pir P, Christen S, Sauer U, Oliver SG, et al. A prototrophic deletion mutant collection for yeast metabolomics and systems biology. Nat Biotechnol. 2012;30: 1176–1178.

52. Peter J, De Chiara M, Friedrich A, Yue J-X, Pflieger D, Bergström A, et al. Genome evolution across 1,011 Saccharomyces cerevisiae isolates. Nature. 2018;556: 339–344.

53. Kahm M, Hasenbrink G, Lichtenberg-Fraté H, Ludwig J, Kschischo M. grofit: Fitting Biological Growth Curves withR. J Stat Softw. 2010;33: 1–21.

54. Wang Y, Backman TWH, Horan K, Girke T. fmcsR: mismatch tolerant maximum common substructure searching in R. Bioinformatics. 2013;29: 2792–2794.

55. Kamrad S, Rodríguez-López M, Cotobal C, Correia-Melo C, Ralser M, Bähler J. Pyphe, a python toolbox for assessing microbial growth and cell viability in high-throughput colony screens. Elife. 2020;9. doi:10.7554/eLife.55160

56. Hartl J, Kiefer P, Meyer F, Vorholt JA. Longevity of major coenzymes allows minimal de novo synthesis in microorganisms. Nat Microbiol. 2017;2: 17073.

57. Decosterd LA, Rochat B, Pesse B, Mercier T, Tissot F, Widmer N, et al. Multiplex ultra-performance liquid chromatography-tandem mass spectrometry method for simultaneous quantification in human plasma of fluconazole, itraconazole, hydroxyitraconazole, posaconazole, voriconazole, voriconazole-N-oxide, anidulafungin, and caspofungin. Antimicrob Agents Chemother. 2010;54: 5303–5315.

58. Mülleder M, Calvani E, Alam MT, Wang RK, Eckerstorfer F, Zelezniak A, et al. Functional Metabolomics Describes the Yeast Biosynthetic Regulome. Cell. 2016;167: 553–565.e12.

59. Jakobson CM, Hartl J, Trébulle P, Mülleder M, Jarosz DF, Ralser M. A genome-to-proteome map reveals how natural variants drive proteome diversity and shape fitness. Science. 2025;390: eadu3198.

60. Demichev V, Messner CB, Vernardis SI, Lilley KS, Ralser M. DIA-NN: neural networks and interference correction enable deep proteome coverage in high throughput. Nat Methods. 2019;17: 41–44.

