## Supplementary Information for "Human metabolites modulate antifungal efficacy and reveal creatinine-mediated antagonism of flucytosine"

Hartl J et al.

Supplementary Figures 1-5
Supplementary References

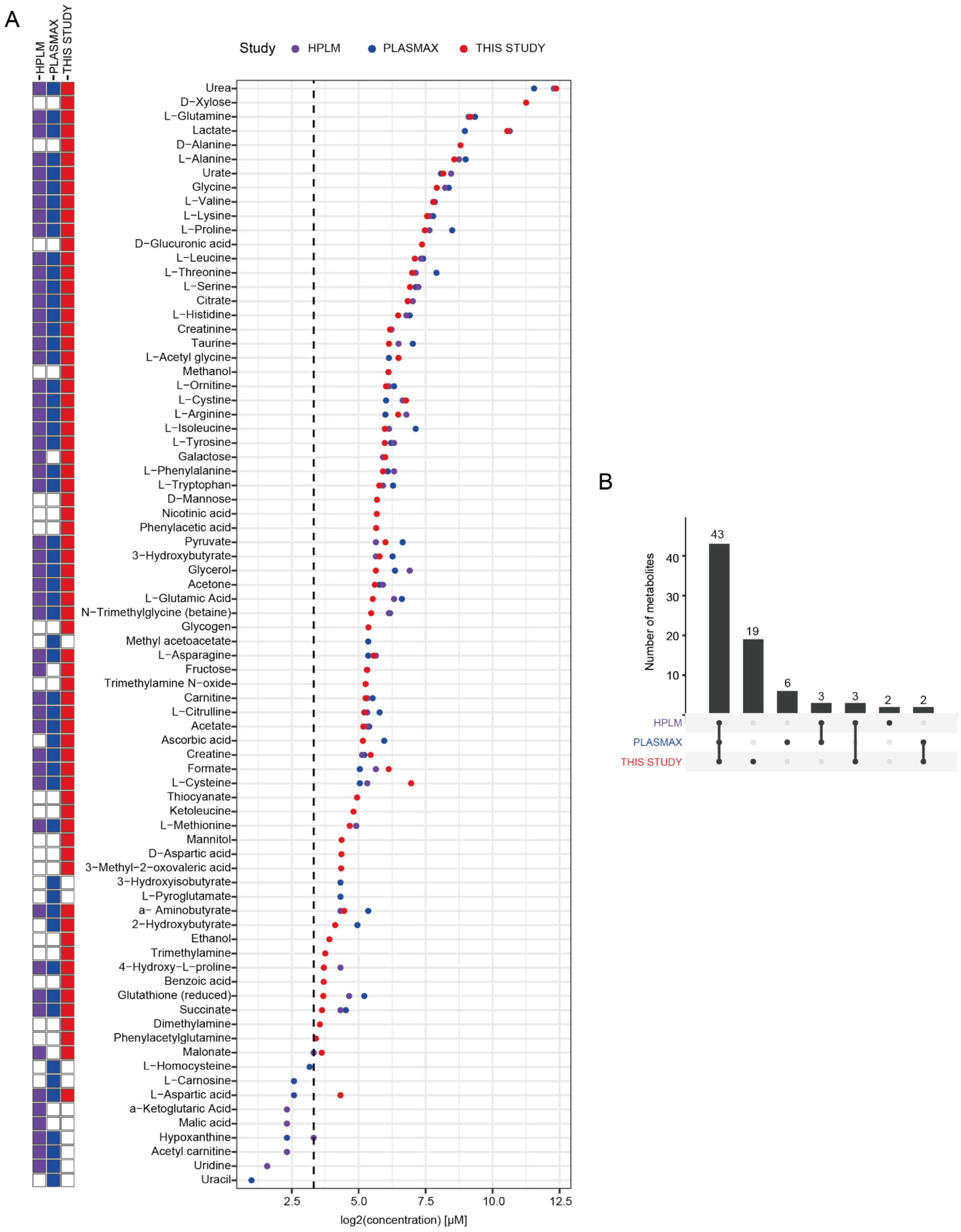


**Supplementary Figure 1. (A)** Comparison of estimated plasma metabolite concentrations from this study (red), human plasma-like medium HPLM (purple) and Plasmax (blue) [1,2]. Latest values for HPLM were taken from https://www.thermofisher.com/de/de/home/technical-resources/media-formulation.360.html (Access 13.05.2026). Boxes to the left indicate presence (colored) or absence (white) of the compound in the respective study. Note that metabolites and molecules used as base formulations of growth media (such as phenol red, glucose as the primary carbon source, essential vitamins, etc. were not included in the analysis). **(B)** Upset plot highlighting intersections of metabolites shown in (A) across studies.

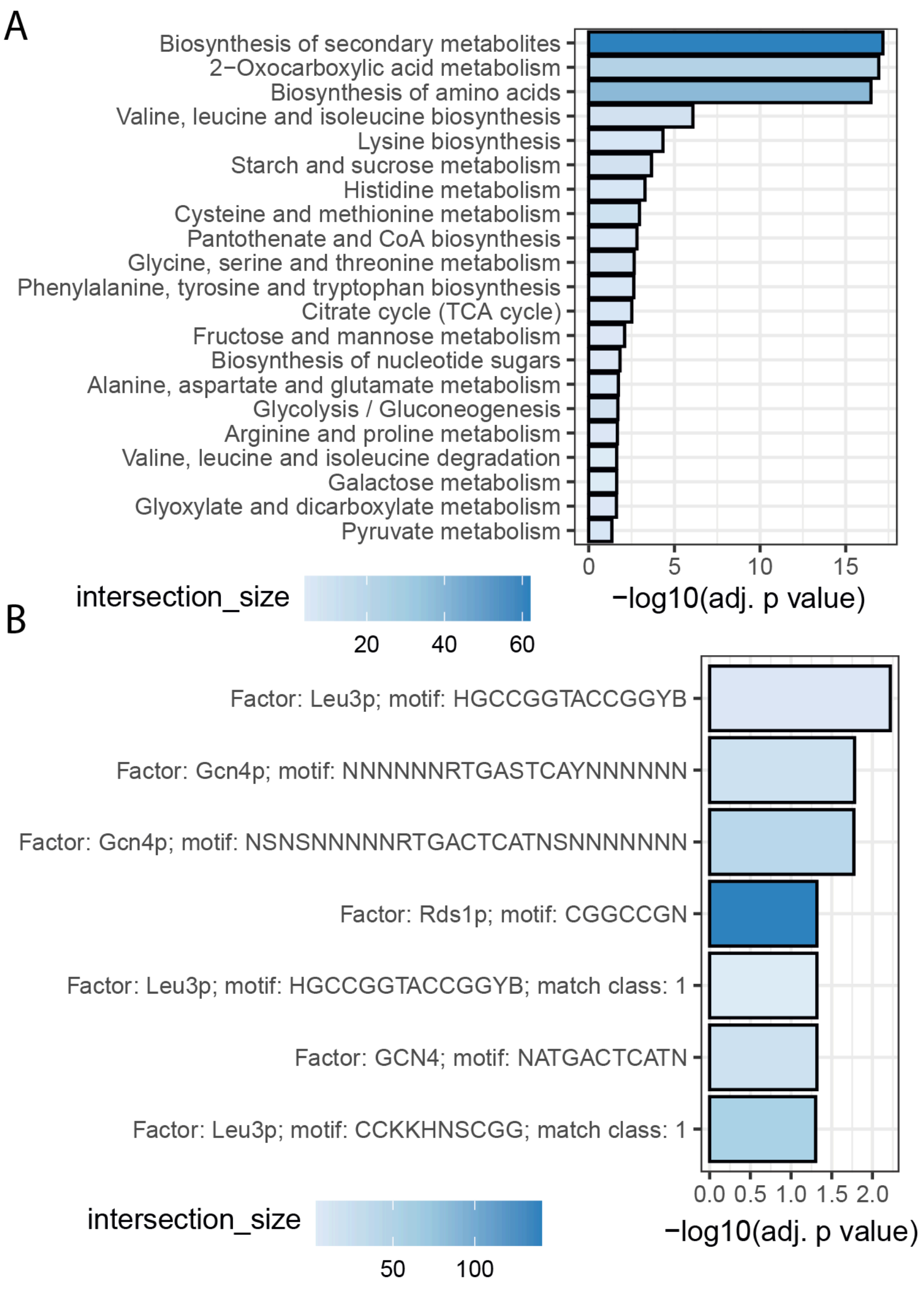


**Supplementary Figure 2.** Enrichments of differentially expressed proteins between *S. cerevisiae* in minimal media and supplemented with human serum metabolite mix. For details, see methods. Briefly, differential protein abundance between SM and SM+HBMM was assessed, and enrichment analysis was performed on significantly changing proteins using gprofiler2 with FDR correction (0.05 threshold), using all quantified proteins as the background. Shown are (**A**) KEGG pathways terms and (**B**) enriched transcription factor motifs. Terms considered redundant were manually removed before visualization.

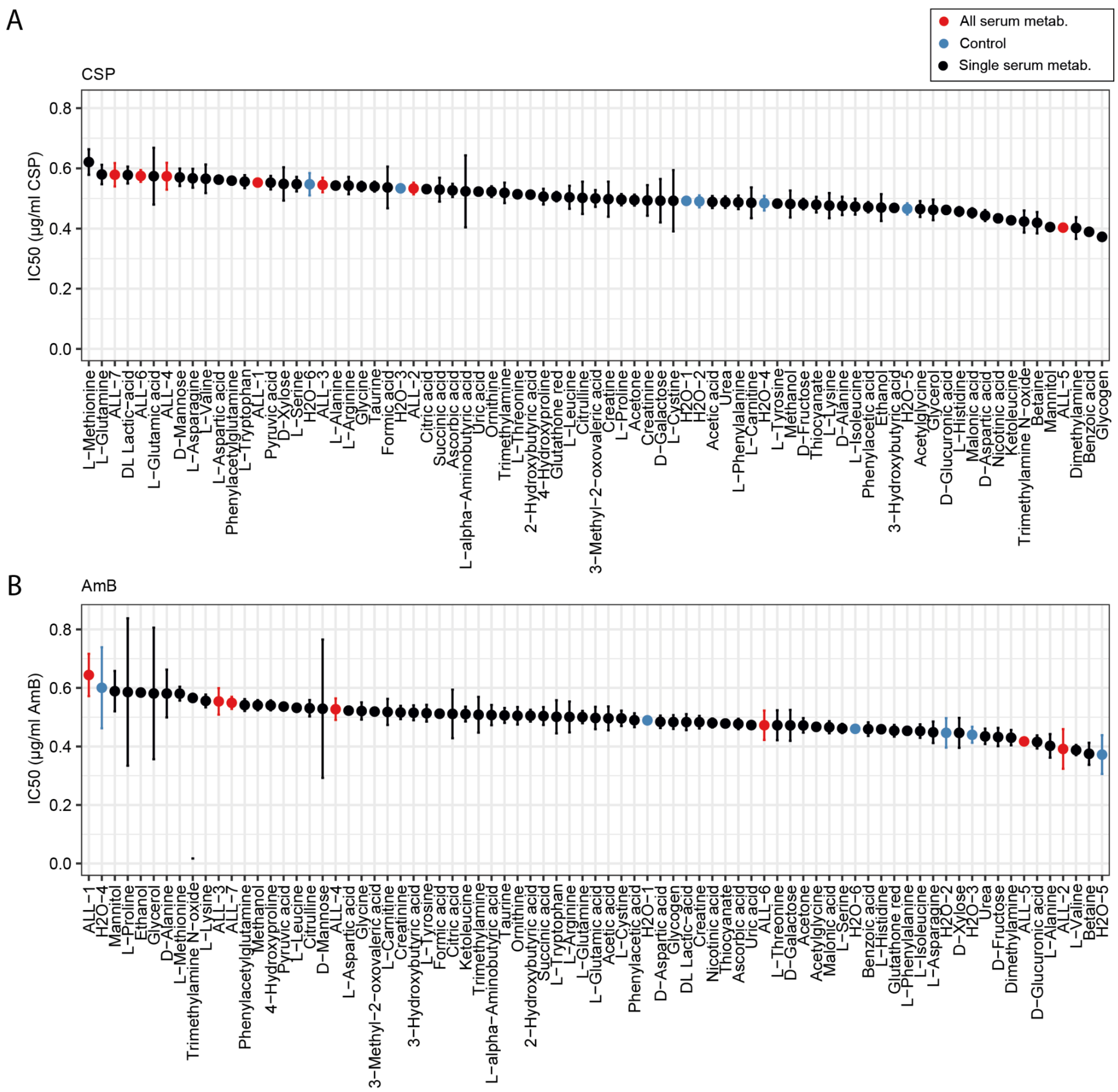


**Supplementary Figure 3.** No substantial antifungal–metabolite interactions for caspofungin and amphotericin B. Data from the antifungal–metabolite interaction screen, in which the sensitivity of *S. cerevisiae* to an antifungal in SM alone, or in the presence of either no (blue), all (red) or individual (black) blood metabolites at physiological concentrations was assessed by microbroth dilution assays using six concentrations of antifungal and two untreated controls. $IC_{50}$ values were estimated from dose–response curves based on OD600. Screening results generated as described for the main screen are shown for caspofungin (A) and amphotericin B (B). $IC_{50}$ values after 24 h of growth are shown; error bars indicate the lower and upper bounds of the 95% confidence interval derived from the dose–response model.

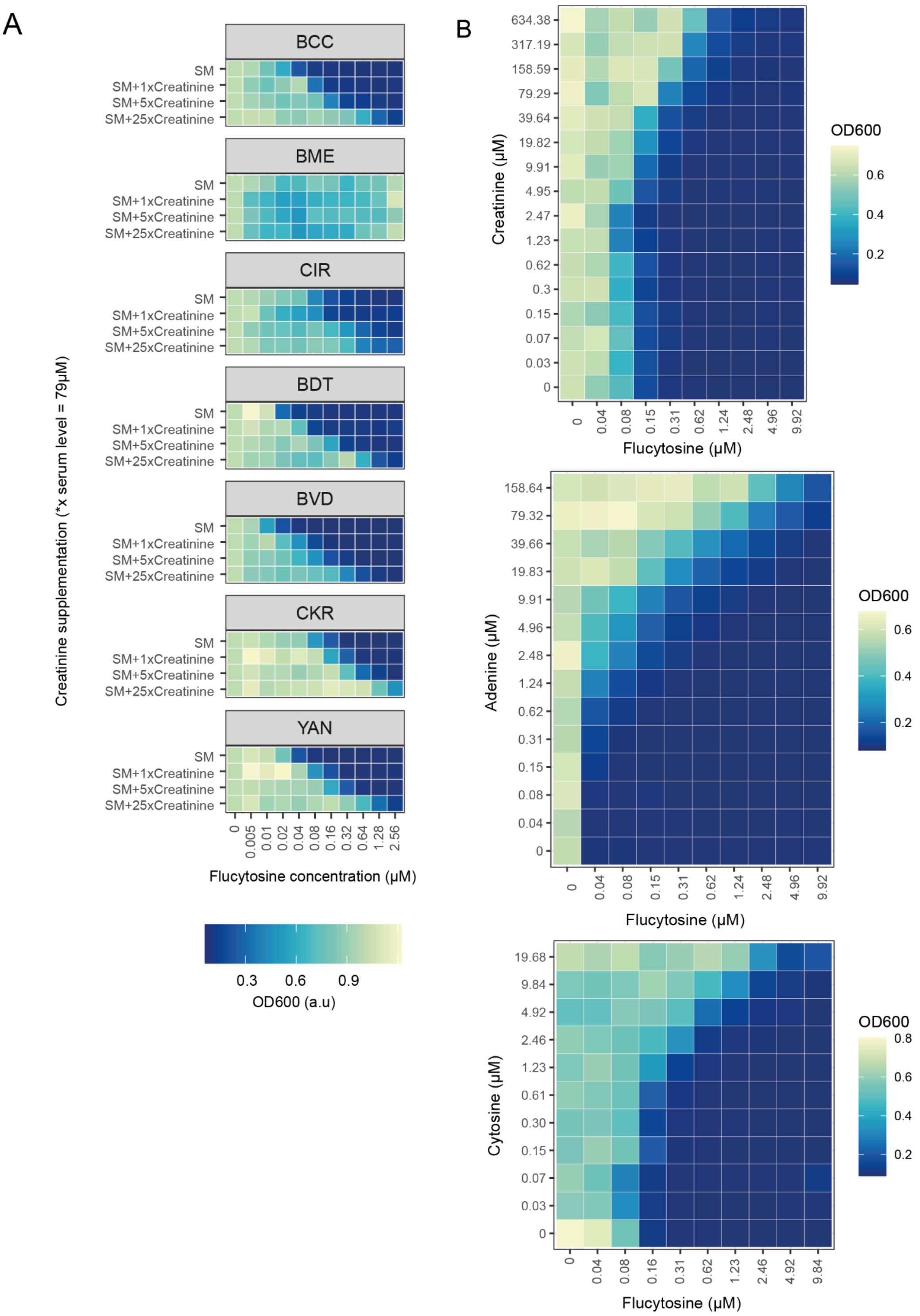


**Supplementary Figure 4.** (**A**) A set of *S. cerevisiae* isolates from the 1011 genome project [3] grown in synthetic minimal (SM) media, or with the addition of increasing levels of creatinine relative to the estimated concentration in human serum (1x; 79 µM, 5x; 395 µM, 25x; 1975 µM) against increasing concentrations of flucytosine (as indicated on the x-axis). Heatmap shows $OD_{600}$ as determined after 24 hours of incubation. For all susceptible isolates, creatinine supplementation led to reduced flucytosine efficacy. (**B**) Checkerboard-like assay comparing growth of *S. cerevisiae* cells ($OD_{600}$ after 30 hours) under different concentration-regimes of flucytosine (x-axis) against creatinine (top), adenine (middle), and cytosine (bottom) concentration (y-axis), respectively. Note that the concentration of metabolites differ, as cytosine has the strongest antagonistic effect against flucytosine, followed by adenine, and creatinine.

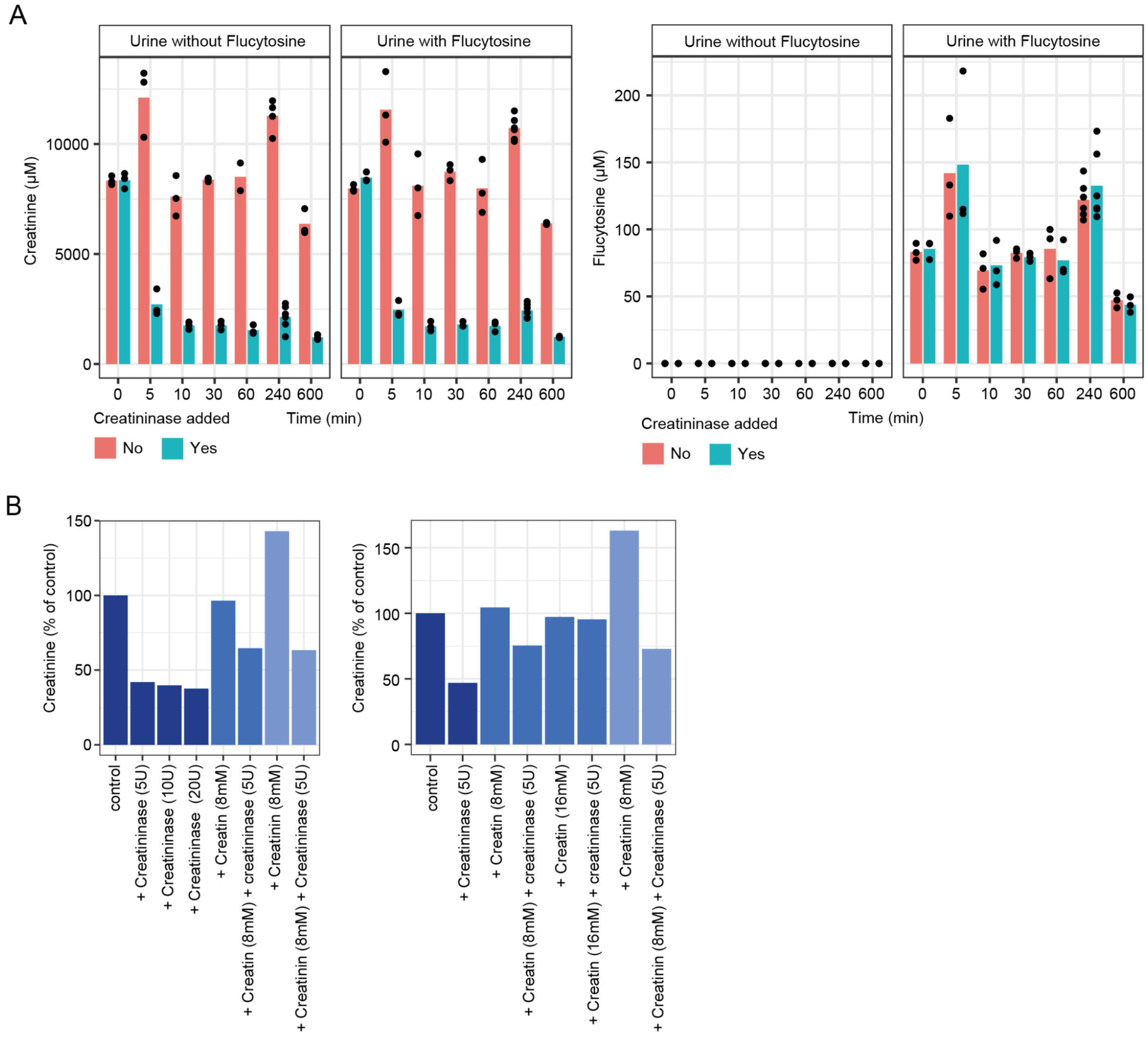


**Supplementary Figure 5.** Creatininase reduces creatinine levels in urine without affecting flucytosine levels. (**A**) Left panel: Creatinine concentration as determined by LC-MS/MS in human urine (left), or human urine with the addition of 150 µM flucytosine (right) at different timepoints after incubation at 37 °C. Where indicated, creatininase was added (turquoise), rapidly reducing creatinine levels. Right panel: as on the left, but showing flucytosine concentration as determined by LC-MS/MS. Note that creatininase addition has no effect on flucytosine levels. (**B**) Creatinine levels in human urine after 1h incubation at 37 °C with the addition of varying amounts of creatininase, creatine, and creatinine. Additional creatininase enzyme has no strong effect, but extra creatine or creatinine led to increase in residual creatinine after creatininase treatment. Each bar represents n=1 replicate per condition from two independent experiments.

**Supplementary references**


1. Cantor JR, Abu-Remaileh M, Kanarek N, Freinkman E, Gao X, Louissaint A Jr, et al. Physiologic Medium Rewires Cellular Metabolism and Reveals Uric Acid as an Endogenous Inhibitor of UMP Synthase. Cell. 2017;169: 258–272.e17.

2. Vande Voorde J, Ackermann T, Pfetzer N, Sumpton D, Mackay G, Kalna G, et al. Improving the metabolic fidelity of cancer models with a physiological cell culture medium. Sci Adv. 2019;5: eaau7314.

3. Peter J, De Chiara M, Friedrich A, Yue J-X, Pflieger D, Bergström A, et al. Genome evolution across 1,011 Saccharomyces cerevisiae isolates. Nature. 2018;556: 339–344.
